# Sporadic phage defense in epidemic *Vibrio cholerae* mediated by the toxin-antitoxin system DarTG is countered by a phage-encoded antitoxin mimic

**DOI:** 10.1101/2023.12.14.571748

**Authors:** Kishen M. Patel, Kimberley D. Seed

## Abstract

Bacteria and their viral predators (phages) are constantly evolving to subvert one another. Many bacterial immune systems that inhibit phages are encoded on mobile genetic elements that can be horizontally transmitted to diverse bacteria. Despite the pervasive appearance of immune systems in bacteria, it is not often known if these immune systems function against phages that the host encounters in nature. Additionally, there are limited examples demonstrating how these phages counter-adapt to such immune systems. Here, we identify clinical isolates of the global pathogen *Vibrio cholerae* harboring a novel genetic element encoding the bacterial immune system DarTG and reveal the immune system’s impact on the co-circulating lytic phage ICP1. We show that DarTG inhibits ICP1 genome replication, thus preventing ICP1 plaquing. We further characterize the conflict between DarTG-mediated defense and ICP1 by identifying an ICP1-encoded protein that counters DarTG and allows ICP1 progeny production. Finally, we identify this protein as a functional antitoxin that abrogates the toxin DarT likely through direct interactions. Following the detection of the DarTG system in clinical *V. cholerae* isolates, we observed a rise in ICP1 isolates with the functional antitoxin. These data highlight the use of surveillance of *V. cholerae* and its lytic phages to understand the co-evolutionary arms race between bacteria and their phages in nature.

**Importance:** The global bacterial pathogen *Vibrio cholerae* causes an estimated 1 to 4 million cases of cholera each year. Thus, studying the factors that influence its persistence as a pathogen is of great importance. One such influence is the lytic phage ICP1, as once infected by ICP1, *V. cholerae* is destroyed. To date, we have observed that the phage ICP1 shapes *V. cholerae* evolution through the flux of anti-phage bacterial immune systems. Here, we probe clinical *V. cholerae* isolates for novel anti-phage immune systems that can inhibit ICP1 and discover the toxin-antitoxin system DarTG as a potent inhibitor. Our results underscore the importance of *V. cholerae* and ICP1 surveillance to elaborate novel means by which *V. cholerae* can persist in both the human host and aquatic reservoir in the face of ICP1.

## Introduction

Bacteriophages (phages) and their bacterial hosts demonstrate real-time co-evolution. At the interface of their conflicts are (typically mobile) genetic elements in the bacterial genome that contain defense systems to subvert phage attacks. Once a phage adsorbs to the host cell and injects its genome, bacteria defend themselves with a variety of systems whose actions can be simplified into two major strategies: mechanisms that (1) target and destroy the phage genome early enough to prevent host takeover and cell death (cell-autonomous immunity) and (2) protect the bacterial population through the death of the infected cell prior to the completion of the phage lifecycle (abortive infection)^1^. Defense systems that mediate cell-autonomous immunity are widespread in bacteria, with restriction-modification (R-M) systems being found in over 75% of bacterial genomes and CRISPR-Cas systems in 40% of genomes^2,3^. Similarly, systems with abortive infection phenotypes are found across many bacteria. For example, retrons, CBASS, and AbiEII are found in 10 to 17% of prokaryotic genomes^4^. Bacterial strains often encode multiple defense systems against phages at both the single cell and population level^4^. Here, having a single cell with a diverse anti-phage arsenal offers an evolutionary advantage against the many phages that could infect the cell. If a phage evolves counter-defense mechanisms to these bacterial defenses, the single cell will be successfully infected, and the phage will propagate. However, if the bacterial population has a diverse, heterogeneous anti-phage system repertoire, the population will resist total collapse in the face of phage with varying counter-defenses, a concept otherwise known as “pan-immunity”^1^. As both bacteria and phage co-evolve, phage-driven loss and gain of bacterial immune systems from bacterial populations are expected; however, studies addressing naturally occurring bacteria-phage pairings are rare, and thus, the field has yet to capture these expected fluctuations fully.

Toxin-antitoxin (TA) systems, two-part complexes consisting of a toxin whose activity leads to bacteriostatic or bactericidal activities and an antitoxin that inhibits the toxin, can underlie abortive infection phenotypes and are ubiquitous in bacterial genomes^5^. Though the direct involvement of TA systems in phage defense is still disputed^2,6^, a growing body of work has shown that several TA systems limit viral spread^7–10^. Dramatic perturbations to cellular physiology, like what unfolds during phage infection, can disrupt the equilibrium between toxin and antitoxin. In these scenarios, the liberated toxin, now free from antitoxin antagonism, abrogates phage production. How the toxin limits phage progeny production in the cell varies: the TA system may directly inhibit the phage lifecycle and/or inhibit the host bacterium by targeting essential cellular processes, thus preventing phage progeny production^11^. Despite mechanistic variability in how defense systems, like TA systems or others, function, there are some commonalities among all, including the tendency for defenses to be encoded in genetic loci containing other defense systems and to be encoded on putatively mobile genetic elements^12–14^. Many anti-phage TA systems reported to date have been studied outside of their native genomic context; thus, investigating natively encoded TA systems near other anti-phage systems could expand our understanding of how defense systems function and how phages overcome such defenses.

One model system to study the dynamics between phage and bacteria is with the bacterium *Vibrio cholerae*, the causative agent of the diarrheal disease cholera, and the lytic bacteriophage ICP1. In both aquatic reservoirs and the human gut, *V. cholerae* faces ongoing predation by ICP1^15,16^. This phage has been co-isolated with *V. cholerae* from cholera patient stool samples from 2001 to the present^17^. Surveillance of *V. cholerae* and ICP1 has provided novel insights into the evolutionary influence that phages and bacteria exert on one another in nature. Clinical isolates of *V. cholerae* can carry mobile genetic elements (MGEs) that defend against invading DNA from plasmids or phages^18–21^. However, only two of these MGEs, the integrative and conjugative elements (ICEs) of the SXT/R391 family and the phage inducible chromosomal island-like elements, PLEs, have been shown to block ICP1, suggesting that these two families of MGEs drive ICP1 resistance in clinical *V. cholerae* strains. SXT ICEs encoded by *V. cholerae* contain an anti-phage locus harboring variable defense systems. Specifically, the two most common SXT ICEs found in *V. cholerae, Vch*Ind5 and *Vch*Ind6, restrict phage by encoding a bacteriophage exclusion (BREX) system or multiple R-M systems, respectfully. Both SXT ICEs function against multiple clinically relevant lytic vibriophages^20^. The PLE, in contrast, is a viral satellite that provides specific protection against ICP1. During infection of a PLE-containing host with ICP1, PLE interferes with ICP1’s lifecycle^22^ while simultaneously hijacking ICP1 proteins^23,24^ to spread itself via transduction^25^. The outcome of PLE activity leads to the complete inhibition of ICP1 progeny in addition to successful PLE mobilization and integration into the surrounding population of cells. To date, 10 PLEs have been discovered^26^. All PLEs studied restrict ICP1; as such, PLE can be considered an anti-ICP1 MGE, and its persistence and evolution over the last century speaks to the longstanding conflict between *V. cholerae* and ICP1 in nature.

To date, 67 ICP1 isolates collected over a >20 year period have been sequenced, which informs conserved genes comprising ICP1’s core genome and fluctuating genomic content that often encodes counter-defenses^17^. To both known defense elements in *V. cholerae*, counter adaptations in ICP1 have been discovered. ICP1 encodes an anti-BREX protein, OrbA, to overcome *V. cholerae* isolates that harbor the SXT ICE *Vch*Ind5. To overcome the R-M-based defense encoded by SXT ICE *Vch*Ind6, ICP1 can gain epigenetic modifications^20^. ICP1 has evolved several sequence-specific counter-defenses to overcome PLEs, including a CRISPR-Cas system to target the PLE genome^27–29^. Interestingly, the prevalence of both PLEs and SXT ICEs in clinical *V. cholerae* isolates varies over time. For example, some PLEs persist in clinical isolates of *V. cholerae* over many endemic waves, whereas others only appear sporadically^26^. ICP1 counteradaptations to the PLE or SXT ICE are expected to contribute to the observed dynamics of varying defenses in epidemic strains of *V. cholerae*^20,26^; however, the potential role that additional defense systems play in limiting the selective advantage of fluctuating defense elements in *V. cholerae* has not been explored.

Given what is known about both PLE and SXT ICE-mediated defense, clinical *V. cholerae* isolates can be expected to be productively infected by ICP1 phages with naturally occurring counter-defenses against the two. However, if clinical *V. cholerae* isolates do not show the expected phage sensitivity pattern, it could suggest novel variants of a previously known defense system or the gain of a new defense system entirely. Here, we discovered a novel anti-phage defense element in clinical isolates of *V. cholerae* from 2008 to 2009. We find that this phage defense element (PDE) encodes the anti-phage toxin-antitoxin system DarTG, consisting of the ADP-ribosylase, DarT, and the ADP-deribosylase, DarG^30–32^. We characterized DarTG-mediated ICP1 defense and discovered a previously unstudied core genome-encoded ICP1 protein, Gp145 (named AdfB for anti-DarT factor B), that provides counter-defense against DarTG. Our results demonstrate that additional defense systems influence the warfare between ICP1 and *V. cholerae* and that such defenses may contribute to the observed dynamic fluctuations of known anti-phage MGEs in *V. cholerae*.

## Results

### Identification of a novel anti-phage element in clinical isolates of V. cholerae

While phenotyping phage susceptibility among clinical *V. cholerae* isolates in our collection, a set of strains from 2008 and 2009 caught our attention as their susceptibility patterns did not match known phage resistance determinants considering SXT ICE and PLE alone. The four strains from this period contain PLE3 and the co-circulating anti-phage SXT ICE *VchInd6*^20,26^. Thus, when infecting these strains with ICP1^2011A^, an ICP1 isolate that overcomes PLE3 by encoding a CRISPR-Cas system and overcomes SXT ICE *Vch*Ind6 with genome modifications preventing recognition by the *Vch*Ind6 encoded restriction-modification system, we expected all *V. cholerae* isolates to be sensitive to ICP1 infection (Fig. 1A). However, of the four PLE3^+^/*Vch*Ind6^+^ *V. cholerae* isolates, only one was susceptible to ICP1^2011A^ similar to the engineered PLE3^+^ and SXT ICE *Vch*Ind6^+^ controls (Fig. 1B, Fig S1). This observation suggested that a novel anti-phage feature or phage defense element was circulating in some *V. cholerae* isolates.

**Figure 1:**
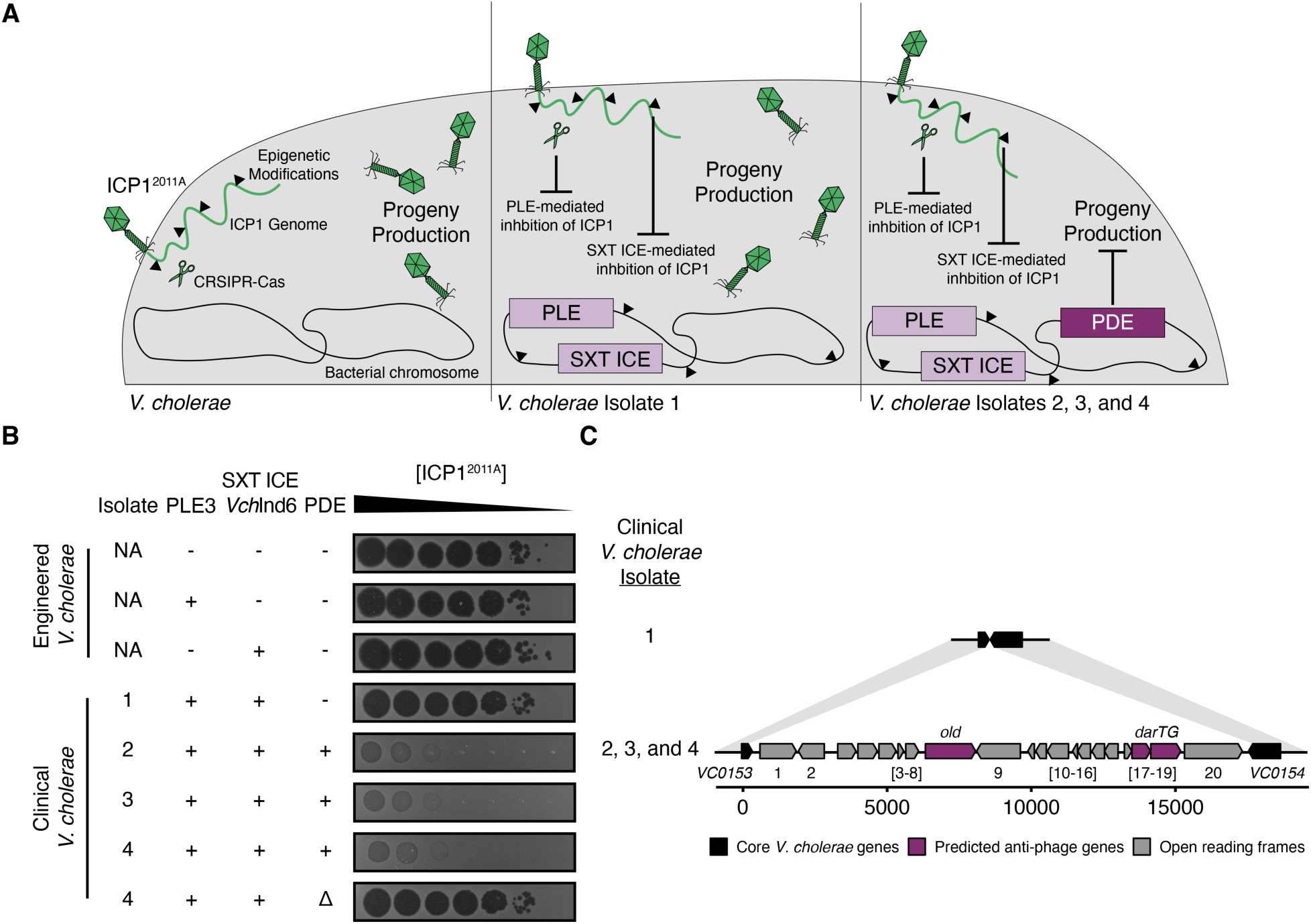
Clinical *V. cholerae* isolates harbor a novel anti-phage genetic element. A) Model representing defense and counter-defense adaptations between *Vibrio cholerae* and ICP1. (Left) *V. cholerae* lacking known phage defenses (PLE and SXT ICE) allows ICP1 to produce progeny during infection, resulting in plaque formation. (Middle) Clinical *V. cholerae* isolates from 2008 and 2009 have multiple defense systems against ICP1, such as PLE3 and the integrative and conjugative element SXT ICE *VchInd6*. ICP1^2011A^ is naturally equipped with the appropriate counter-defense measures (CRISPR-Cas and epigenetic modifications^20,29^) to mount a successful infection of PLE3^+^/*Vch*Ind6^+^ *V. cholerae*. (Right) Multiple clinical *V. cholerae* isolates from 2009 inhibited ICP1^2011A^ plaquing, suggesting they encode another phage defense element (PDE). B) Ten-fold serial dilutions of ICP1^2011A^ were spotted on engineered or clinical *V. cholerae* strains encoding PLE3, SXT ICE *Vch*Ind6, or the novel PDE. Black zones of clearing are where ICP1^2011A^ produced plaques. The opaque background is the *V. cholerae* lawn. Engineered strains are isogenic, while clinical isolates are not. “-”, “+”, “Δ” indicates the absence, presence, and knock-out, respectfully, of the genetic element indicated. These assays are representative of three biological replicates (additional replicates are shown in Fig. S1). C) Genetic map (drawn to scale) of the PDE located between genes *VCA0153* and *VCA0154*. Open reading frames not predicted to defend against phages are grey. The predicted anti-phage systems *old* (nuclease) and *darTG* (toxin-antitoxin system) are colored in purple. Areas of >90 percent nucleotide identity are shaded in light grey.

The most notable genomic difference between the phage-resistant and phage-susceptible *V. cholerae* isolates was a putative phage defense element (PDE) encoding 20 predicted open reading frames (ORFs) (Fig. 1C). Amongst these 20 ORFs, we identified an integrase (an enzyme involved in the mobility of MGEs)^33^, transcriptional regulators, and hypothetical proteins (summarized in Supplemental Data Sheet 1). Most notably, two potential anti-phage systems were identified: ORF8 encoding an AAA+ family ATPase with predicted structural similarity^34^ to an OLD-family nuclease, an enzyme that can prevent DNA replication of bacteriophage lambda in *Escherichia coli*^35,36^ and ORFs18-19, encoding the predicted toxin-antitoxin (TA) system DarTG, a system also shown to inhibit phage genome replication in *E.* coli^31^. Further investigation of the PDE revealed similar elements in other *V. cholerae* strains and *Vibrio* species with nucleotide homology to the integrase and/or ORFs 3-7 and 10-16, which have no known function in phage defense. While most of these PDE-like elements have yet to be characterized, one of them is *Vibrio* Seventh Pandemic Island 2 (VSP-2), which encodes the Lamassu family defense system, DdmABC^18^ (Fig. S2). The high level of nucleotide homology across these elements could promote the exchange of diverse anti-phage “cargo” genes to provide pan-immunity.

To confirm that the PDE was indeed the novel genomic feature responsible for phage-resistance in these clinical isolates, we pursued deleting the element in the clinical *V. cholerae* isolate four as a representative isolate. However, we were not initially successful in generating a deletion mutant of the entire element by homologous recombination. As TA systems have also been implicated in genetic element maintenance^6^, we reasoned that the DarTG toxin-antitoxin system might limit our capacity to delete the PDE. Therefore, we first deleted *darT,* encoding the toxin, before attempting to delete the full element. The transformation efficiency of the *ΔdarT* strain was improved compared to the wild type, indicating that this toxin-antitoxin pair contributes to the maintenance of the element (Fig. S3). Upon deleting the PDE from the clinical *V. cholerae* isolate, we observed the restoration of plaquing by ICP1^2011A^ (Fig. 1B, Fig S1), indicating that this novel element is the necessary component for phage inhibition in these clinical isolates.

To determine if the *V. cholerae* PDE is sufficient to restrict phage, we introduced the element into its natural integration locus via natural transformation in our permissive laboratory *V. cholerae* strain (herein referred to as PDE^+^ *V. cholerae*). This strain lacks other anti-phage elements like the SXT ICE and PLE. We then challenged the PDE^+^ *V. cholerae* strain with the most extensively characterized ICP1 isolate^17,22,37^, ICP1^2006E^, and observed the inhibition of plaque formation similar to that of ICP1^2011A^ (Fig. S4). These results indicate that the PDE in some clinical isolates of *V. cholerae* is sufficient to restrict genetically distinct ICP1 isolates. To understand if this PDE inhibits other vibriophages, we challenged the engineered strain with ICP2 and ICP3, podoviruses that are unrelated to each other and ICP1^15^. Here, we saw a reduction in plaque size of ICP2 and no inhibition of ICP3, indicating that the element does not broadly protect against all vibriophages found in cholera patient stool (Fig. S4). In summary, these data reveal that a novel phage defense element present in some clinical *V. cholerae* isolates is capable of restricting ICP1 plaque formation.

### DarTG is the principal anti-phage system encoded by the V. cholerae PDE

Having established that the PDE is both necessary and sufficient to restrict ICP1 plaquing, we sought to decipher which PDE-encoded genes were responsible. To interrogate if either the OLD-family nuclease homolog (herein *old)* or DarTG is necessary for ICP1 restriction, we deleted both *old* and *darT* separately in PDE^+^ *V. cholerae* and challenged the resulting strains with ICP1^2006E^. We observed inhibition of plaque formation when *old* was deleted, but restoration of plaquing to an efficiency of plaquing (EOP) of one when *darT* was deleted (Fig. 2A). From these results, we concluded that DarTG is the PDE-encoded phage defense system responsible for ICP1 inhibition.

**Figure 2:**
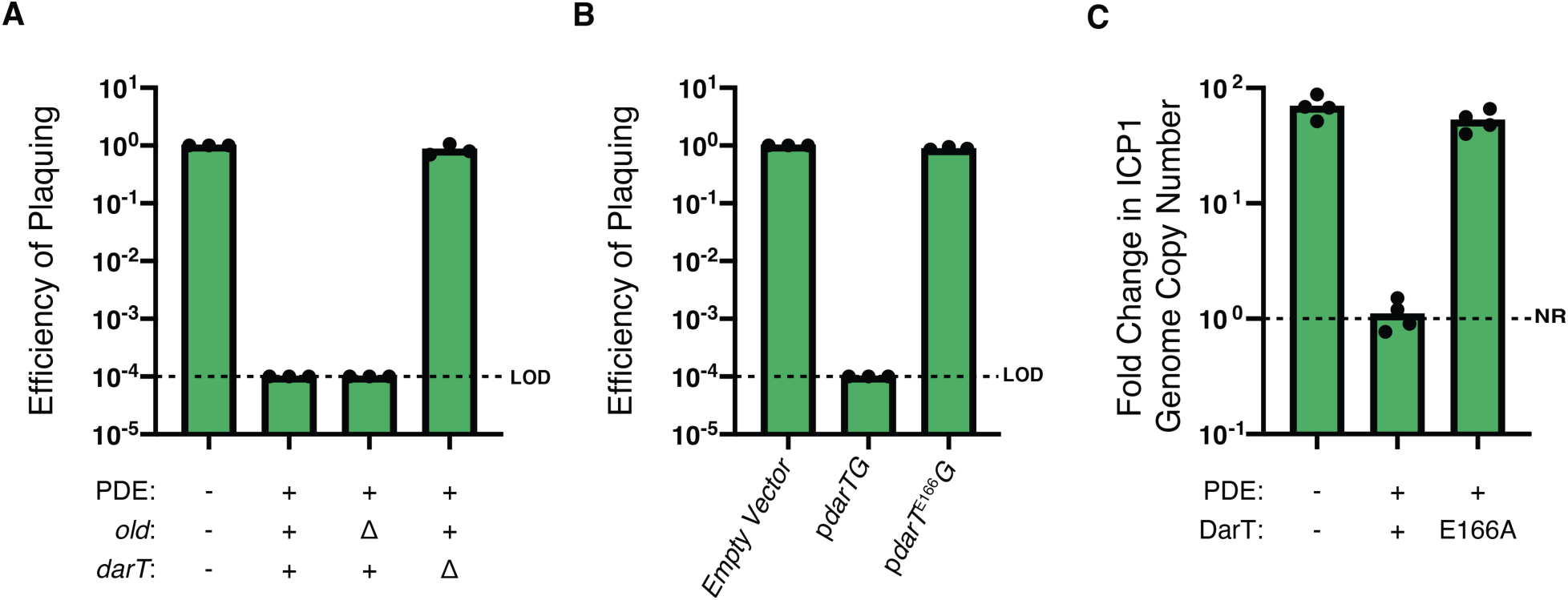
The DarTG system inhibits ICP1^2006E^ plaque formation and genome replication. A) The Efficiency of Plaquing (EOP) of ICP1^2006E^ in PDE^-/+^ *V. cholerae* and its mutant derivatives. For A-B, LOD represents the limit of detection for the assay. For A-C, each dot represents an individual biological replicate. The bar indicates the mean of biological replicates. B) EOP of ICP1^2006E^ in the presence of ectopically expressed DarTG, DarT^E166A^G, or an empty vector control in *V. cholerae*. C) ICP1^2006E^ genome replication after infection of PDE^+^ *V. cholerae* and its mutant derivatives as determined by qPCR. To determine fold change, samples taken 20 minutes post-infection were compared to the input sampled immediately after adding phage. NR represents no replication.

The DarTG system has been shown to inhibit phage infection in *E. coli*^31^. The proposed model for how DarTG inhibits phage is as follows: First, the toxin, DarT, and the antitoxin, DarG, are constitutively expressed in the cell. In this balanced state, DarG prevents DarT activity through steric interactions^38^ and by reversing the catalytic activity of DarT^30^. Through an unknown mechanism, phage infection upsets the equilibrium between DarTG and DarT is freed to add ADP-ribose groups to single-stranded bacterial and phage DNA thus leading to phage inhibition and an abortive infection phenotype^31^. In previous work, two DarTG systems were reported to be sufficient to inhibit *E. coli* phages outside the context of their respective genetic elements^31^. To interrogate if DarTG encoded by *V. cholerae* is sufficient to inhibit ICP1 outside of the context of the PDE, we engineered a plasmid expressing DarTG and introduced it into permissive PDE^-^ *V. cholerae*. As a control, we engineered an inactive DarTG plasmid in which the ADP-ribosylase, DarT, had no catalytic activity (Fig. S5) and introduced it into PDE^-^ *V. cholerae* (pDarT^E166A^G). In the presence of wild type DarTG, we observed that ICP1^2006E^ plaque formation was restricted, indicating that the system is sufficient for protection. Conversely, phage infection was unaffected by the presence of the catalytically inactive DarT, thus demonstrating that phage restriction is dependent on the enzymatic activity of DarT (Fig. 2B).

In *E. coli,* the DarTG system blocks phage genome replication^31^. Therefore, to investigate if the *V. cholerae* DarTG system restricts ICP1 genome replication in its native context, we calculated the fold change in phage genome copy number after a single round of infection of PDE^+^ *V. cholerae,* the permissive PDE^-^ strain or a newly engineered PDE^+^ strain expressing the catalytically dead mutant of DarT. Here, we observed robust phage genome replication during infection of the permissive host and the host with catalytically inactive DarT, while genome replication was not detected following infection of PDE^+^ *V. cholerae* possessing the catalytically active, wild type DarT (Fig. 2C). From these data, we conclude that the catalytic activity of the DarT is necessary to inhibit phage genome replication and subsequent plaque formation.

### ICP1 escapes DarTG-mediated inhibition through a nonsynonymous mutation in gp145

Phages can bypass toxin-antitoxin systems through mutations that allow the phage to avoid the toxin’s activity^31^, by encoding an inhibitor of the system^31,39–42^, or by mutation of the trigger of the system^10^. To probe if ICP1 can evolve to overcome *V. cholerae*-encoded DarTG, we selected for escape phages by infecting the clinical PDE^+^ *V. cholerae* (isolate 4, Fig. 1) with ICP1^2011A^ and picking rare plaques. In total, we isolated three escape phages with improved plaquing efficiency in the presence of the PDE (Fig. S6). We sequenced the escape phages and then compared the genomes to the parental phage using BREseq^43^ (Supplemental Table 1). From the analysis, a single, conserved point mutation resulting in an amino acid change from glycine to aspartic acid at position 62 (G62D) in the hypothetical gene product, Gp145 (Protein ID: AXY82238.1), was identified. This gene product is considered a member of ICP1’s core genome, yet there is no predicted function for it^17^. These results suggest that Gp145 is involved in the DarTG-mediated inhibition of ICP1.

Recently, phage proteins necessary for phage viability have been determined as activators for phage defense systems, including TA systems^10,44^. To interrogate if Gp145 triggers the DarTG TA system, we introduced inducible plasmids expressing either Gp145^G62^ (wild type) or Gp145^D62^ (escape allele) into PDE^+^ *V. cholerae*. These strains were induced for three hours, and we monitored percent growth by plating for colony-forming units (CFU). Given that DarT over-expression kills *V. cholerae* (Fig. S5), we expected to observe a growth defect in the strain that expressed the wild type Gp145^G62^ if Gp145 triggered the DarTG TA system. However, all strains grew equally well compared to the strain containing the empty vector control (Fig. S7). Furthermore, we found that ICP1^2006E^ engineered with a clean deletion of Gp145 remained susceptible to DarTG (Fig. 3A). These results support the conclusion that Gp145 is not the activator of the DarTG system and suggest that the G62D amino acid substitution in Gp145 is necessary for DarTG-escape.

**Figure 3:**
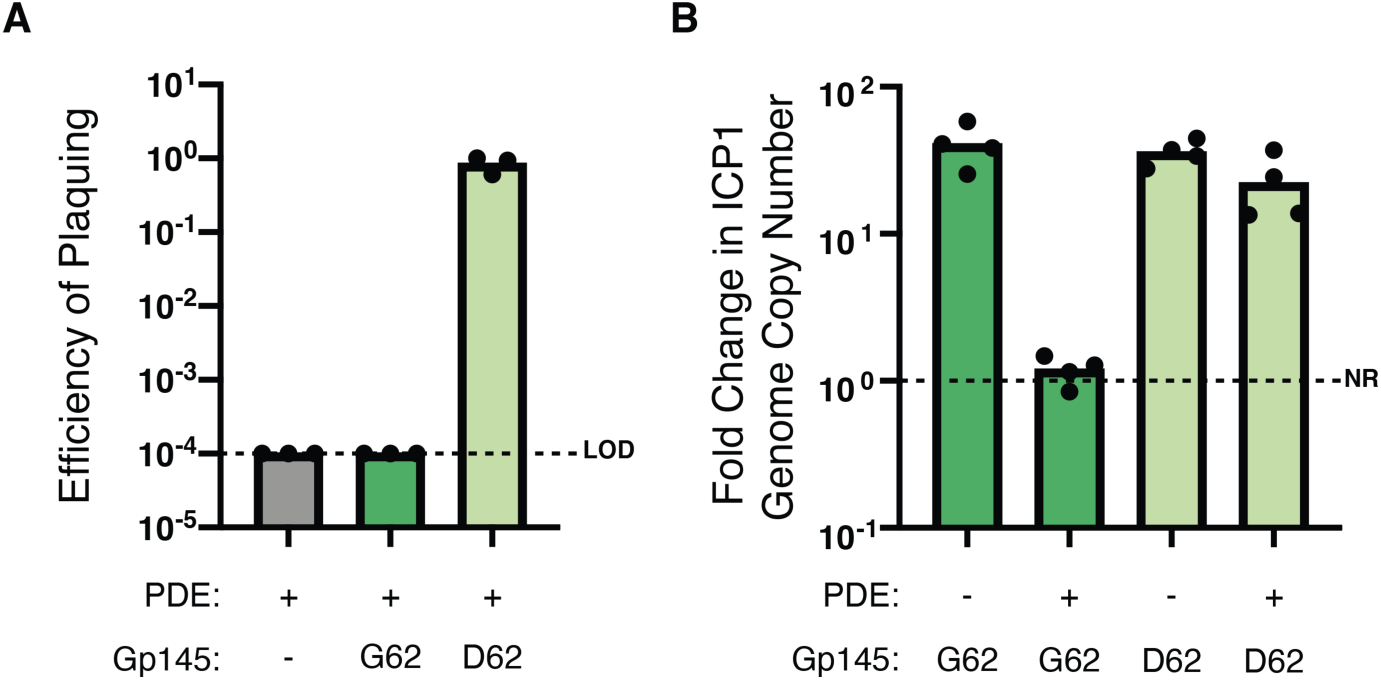
ICP1 escapes DarTG-mediated inhibition through a single point mutation in Gp145 A) The efficiency of plaquing (EOP) of engineered ICP1^2006E^ with the allele of *gp145* indicated on PDE^+^ *V. cholerae*. Each data point is representative of a single biological replicate, while the bar represents the mean of these replicates. LOD is the limit of detection. B) Fold change of genome copy number of isolates of ICP1 engineered to encode either Gp145^G62^ or Gp145^D62^ in the presence or absence of the PDE as determined by qPCR. Fold change was determined by comparing samples 20 minutes post-infection to the input sampled immediately after adding phage. Each data point is representative of a single biological replicate. The bar represents the mean of these replicates. NR represents no replication.

To test the hypothesis that Gp145^D62^ is required to escape the DarTG system, we reconstituted both alleles of Gp145 in ICP1*Δgp145*. When infecting PDE^+^ *V. cholerae* with these newly engineered phages, we observed that Gp145^D62+^ ICP1 could escape the DarTG system (Fig. 3A). Having previously established that DarTG inhibits ICP1 genome replication (Fig. 2C), we next monitored ICP1 genome copy number prior to- and 20 minutes post-infection, hypothesizing that the ICP1 isolate encoding Gp145^D62^ would be able to replicate its genome to the same level as is observed in a permissive host. Indeed, our results show that the mutated allele, Gp145^D62^, allows ICP1 to replicate its genome in the presence of DarTG (Fig. 3B). From these data, we conclude that Gp145^D62^ is necessary for ICP1 to overcome restriction by DarTG.

### The phage-encoded Gp145^D62^ inhibits DarT toxicity in V. cholerae

Gp145^D62^ provides ICP1 robust protection against the DarTG system (Fig. 3A), suggesting it is an inhibitor of DarT activity. To investigate if Gp145^D62^ can abrogate DarT-mediated cell death, we introduced either an empty vector or vectors for the inducible expression of DarG, Gp145^G62^, or Gp145^D62^ into *V. cholerae* with inducible expression of DarT. We then calculated the percent growth of cells by comparing the CFUs pre-and post-expression of DarT and DarG or variants of Gp145 for three hours. Upon expression of DarT and the empty vector control or Gp145^G62^, we observed a reduction in colony formation indicative of cell death (Fig. 4A). However, in the presence of DarT, cells expressing the known antitoxin, DarG, or the escape allele Gp145^D62^, robustly grew (Fig. 4A) demonstrating that Gp145^D62^ abrogates DarT-mediated toxicity to a similar level as the cognate antitoxin DarG.

**Figure 4:**
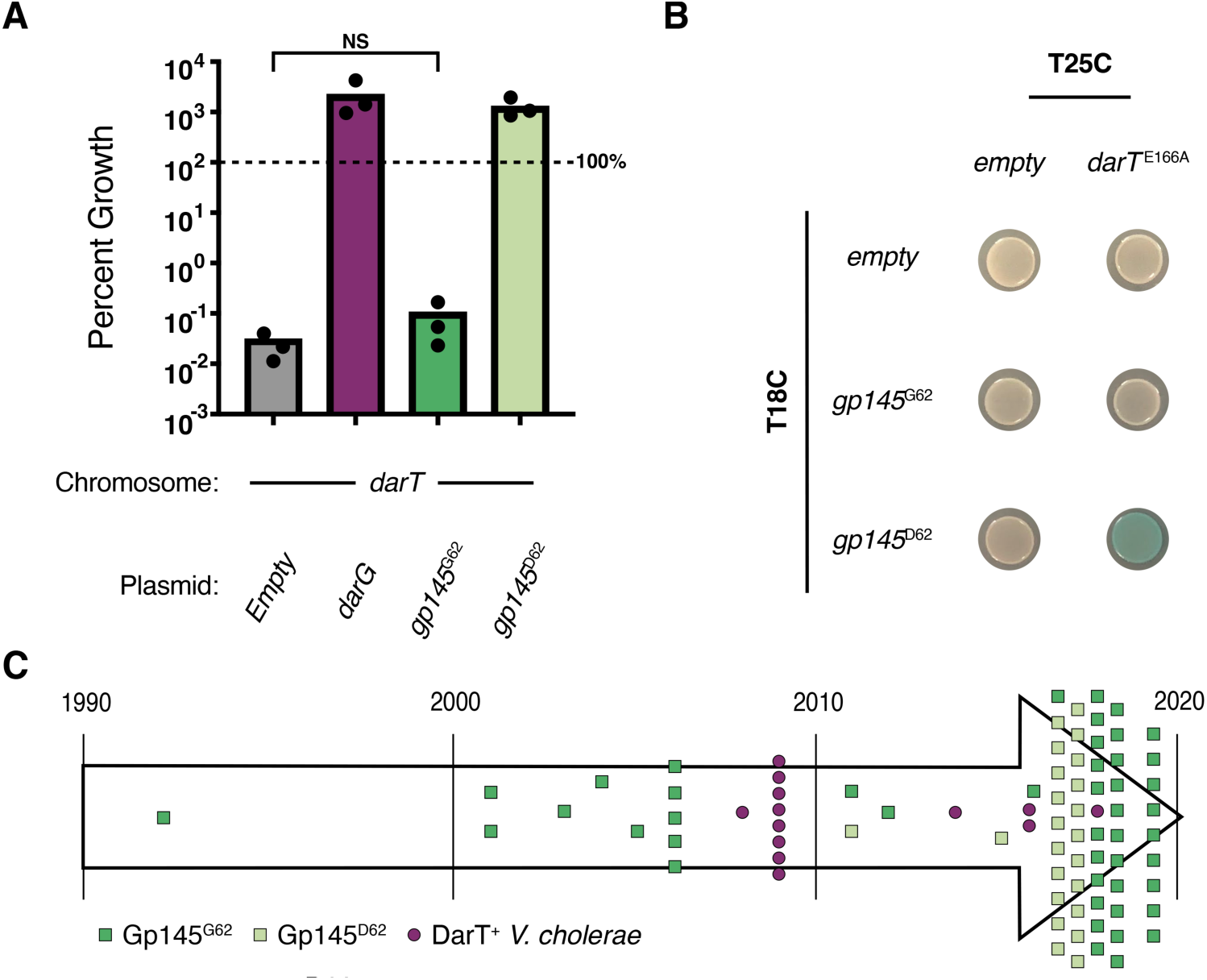
Gp145^D62^ inhibits DarT toxicity in *V. cholerae* and is found in ICP1 in nature. A) Percent growth as determined by colony-forming units of *V. cholerae* cells expressing DarT (chromosomal expression system) along with the indicated vector for three hours. Each dot represents one biological replicate. The bar indicates the mean of all replicates. 100% cell survival is marked by the dotted line. NS represents non-significance by a student’s t-test. B) Bacterial two-hybrid data to detect protein-protein interactions. Cells containing the plasmids T25C-*darT^E^*^166^*^A^* (indicated on top) and T18C-*gp145^G^*^62^ or T18C-*gp145^D^*^62^ (indicated on left) were spotted on agar plates containing X-gal and inducer. Blue spots indicate a physical interaction between the proteins fused to the CyaA subunits. C) Timeline showing isolates of ICP1 and DarT^+^ *V. cholerae* colored according to the legend. Each square represents a genetically distinct ICP1 isolate and is colored according to the *gp145* allele present. Each purple circle represents a genetically distinct DarT containing *V. cholerae* isolate detected in a database of 3,363 *V. cholerae* isolates from 1916 to 2019.

In other systems, DarG abrogates DarT toxicity in two ways: DarG’s N-terminal macrodomain catalyzes the reverse reaction of DarT (ADP-deribosylation), while its C-terminal domain physically occludes the active site of DarT by mimicking negatively charged ssDNA and binding where DarT otherwise would bind to DNA to add ADP-ribose groups^30,38^. To gain insight into the mechanism by which Gp145^D62^ abrogates DarT’s activity, we first compared the amino acid sequence identity between Gp145 and DarG. Because Gp145 is 77 amino acids in length, whereas DarG is 347 amino acids, we chose to compare Gp145 to the truncated N- and C-terminal ends of DarG. In both cases, Gp145 and DarG have low (<15%) sequence similarity. Additional *in silico* analysis by HHPred and I-TASSER^45–47^ of Gp145 did not reveal any predicted enzymatic activity that would counteract DarT-mediated toxicity, suggesting it allows ICP1 to escape DarTG by physically interacting with DarT. This interaction would be similar to the hypothesized mechanism that AdfA (anti-DarT factor A) encoded by <RB69 uses to inhibit DarT in *E. coli*^31^. Gp145 and AdfA share low (<15%) sequence similarity, so for additional insight, we turned to AlphaFold2^48^ to predict the structure of DarG (Fig. S8A) and Gp145^D62^ (Fig. S8B) hypothesizing that Gp145 ^D62^ would be structurally similar to the C-terminus of DarG given that the two inhibit the same DarT toxin. We overlaid the predicted structure of Gp145^D62^ onto the predicted structure of DarG and observed that indeed the C-terminus of DarG and Gp145^D62^ are predicted to be structurally similar despite having low sequence similarity (Fig. S8C). This observation further suggested that Gp145^D62^ interacts with DarT to abrogate toxicity, similar to DarG’s C-terminus^38^. To experimentally determine if Gp145 interacts with DarT, we performed bacterial two-hybrid assays, which use translational fusions of proteins of interest to separate catalytic domains of adenylate cyclase (CyaA). If these fusions come together by protein-protein interactions, β-galactosidase enzyme will be produced, resulting in a positive, blue colony when plated on indicator plates^49^. Consistent with their differential ability to abrogate DarT toxicity, we observed an interaction between DarT and Gp145^D62^ but not Gp145^G62^ (Fig. 4B, Fig. S9). Taken together, these results indicate that Gp145^D62^ functions as an antitoxin mimic by directly binding to abrogate DarT’s activity.

Having established that Gp145^D62^ antagonizes DarT and knowing that Gp145 is present in all isolates of ICP1, we chose to assess the diversity of Gp145 encoded by ICP1. To our surprise, when looking at Gp145 among all 67 genetically distinct ICP1 isolates spanning a collection period between 1992 – 2019^17^, we identified only two naturally occurring variants: Gp145^G62^ and Gp145^D62^. Interestingly, ICP1 isolates possessing Gp145^D62^ succeed the isolation of the three PDE^+^/DarTG^+^ clinical *V. cholerae* isolates from our collection. To better understand the prevalence of DarTG in *V. cholerae* in relation to the circulating *gp145* allele, we expanded our search of DarT found in *V. cholerae* by searching for *darT* in a previously generated database of 3,363 sequenced *V. cholerae* genomes that spans a much greater date and geographic sampling range than our ICP1 collection^26^. Many of these genomes come from the clinical setting. DarT was infrequently detected in *V. cholerae* (13/3,363 genomes); however, all DarT homologs recognized from this search were identified in *V. cholerae* strains after 2008, an observation that coincides with the appearance of the functional antitoxin mimic (Gp145^D62^) in ICP1 isolates (Fig. 4C). Together, the emergence of Gp145^D62^ in ICP1 isolates and the rarity of DarT^+^ *V. cholerae* isolates from the database could suggest the circulation of DarT in under-surveyed environmental *V. cholerae* isolates. Quickly after the appearance of Gp145^D62^ in ICP1 isolates, DarTG^+^ *V. cholerae* isolates in South Asia were no longer detected, potentially suggesting that this counter-defense led to the turnover of DarTG and the PDE from *V. cholerae* populations. Together, these data provide evidence for the natural selection of a Gp145 variant in ICP1 that can abrogate DarTG-mediated anti-phage activity.

After 2018, we see a reversion to Gp145^G62^ that overlaps the loss of DarTG from *V. cholerae* populations (Fig. 4C). Given the return of Gp145^G62^ and the evidence of counter-defense by Gp145^D62^ against DarTG, we hypothesized that Gp145^G62^ may function as an antitoxin to a different DarT toxin not captured in our database. Thus, we expanded our search to understand how the PDE-encoded DarT fits into previously characterized DarT proteins across different bacteria. We built a phylogenetic tree of ∼5000 DarT sequences, including DarT from the PDE^+^ *V. cholerae* (Fig. 5A). Previous bioinformatics analysis of 16 DarT homologs classified the proteins into five groups based on percent identity to the crystalized *Thermus* species DarT^32^. From the tree, we observed that the previous groupings defined by amino acid similarity across DarT homologs remain consistent, except for group 2 DarT proteins (Fig. 5A). Interestingly, DarTs from anti-phage DarTG systems previously described in *E. coli*^31^ and this study are not limited to any one clade, suggesting that diverse DarTG systems can function in phage defense.

**Figure 5:**
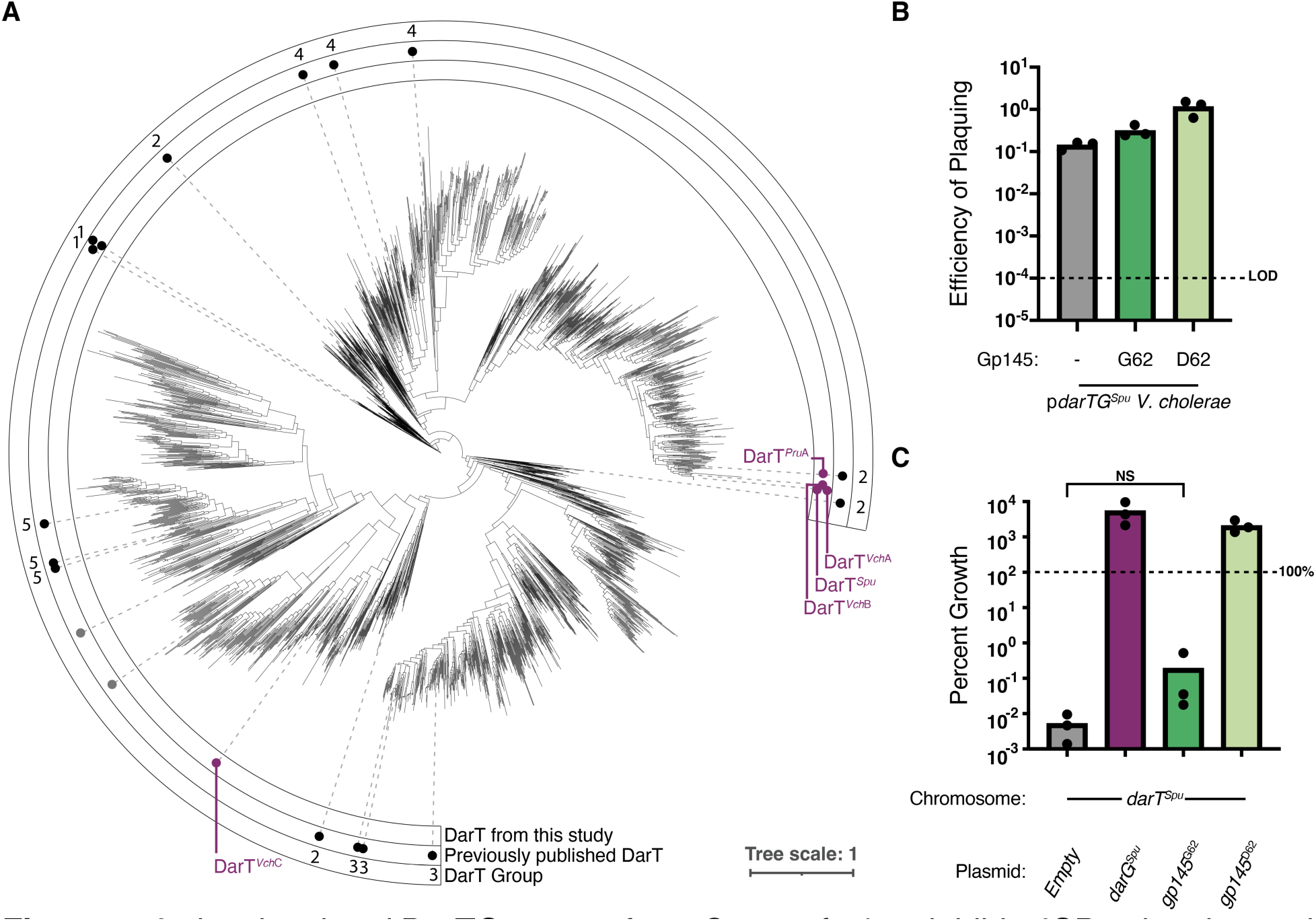
A closely related DarTG system from *S. putrefaciens* inhibits ICP1 plaquing and is also overcome by Gp145^D62^. A) Phylogenetic tree of DarT proteins clustered at a >80% amino acid identity. In the innermost ring, purple dots indicate DarTs from this study, where DarT*^Vc^*^hA^ is the one encoded by the PDE. In the middle ring, black dots are DarT proteins described in Scheller *et al.* (2021)^32^, and grey dots indicate DarT proteins that provide phage defense as described in LeRoux *et al.* (2022)^31^. The numbers in the outer ring indicate which DarT group the previously published DarTs belong to. B) Efficiency of plaquing (EOP) of engineered ICP1^2006E^ with the allele of *gp145* indicated on pDarTG*^Spu^ V. cholerae*. LOD is the limit of detection. C) Percent growth as determined by colony-forming units of *V. cholerae* cells expressing DarT*^Spu^*(chromosomal expression system) along with the vector indicated for three hours. 100% cell survival is marked by the dotted line. Each dot represents one biological replicate. The bar indicates the mean of all replicates. NS represents non-significance by a student’s t-test.

To determine if Gp145^G62^ defended ICP1 against diverse DarTG systems, we used the phylogenetic tree to select other DarTG systems that are similar or distantly related to the DarTG system from clinical *V. cholerae* isolates characterized during this study (designated DarTG*^Vch^*^A^). The sample group included systems from *Shewanella putrefaciens* (DarTG*^Spu^*), *Providencia rustigianni* (DarTG*^Pru^*), and other *V. cholerae* strains (DarTG*^Vch^*^B^ and DarTG*^Vch^*^C^) (Supplemental Table 2). To assess anti-phage activity of these systems, we engineered plasmids expressing these DarTG systems, introduced them into permissive PDE^-^ *V. cholerae*, and infected these strains with ICP1 that lacks Gp145 or encodes either Gp145^G62^ or Gp145^D62^. Most of these systems (DarTG*^Pru^*, DarTG*^Vch^*^B^, and DarTG*^Vch^*^C^) did not inhibit ICP1 plaquing (Fig. S10), precluding us from assessing if phages encoding variants of Gp145 could overcome these systems. However, ICP1 lacking Gp145 was inhibited by the closely related system DarTG*^Spu^* (Fig. 5B). When infecting pDarTG*^Spu^ V. cholerae*, ICP1 encoding Gp145^G62^ showed a subtle increase in EOP, while ICP1 encoding Gp145^D62^ returned to an EOP of 1 (Fig. 5B), suggesting that *gp145*^D62^ is also a gain of function mutation in the face of a different DarTG system. Accordingly, we observed that expression of Gp145^D62^ abrogated the toxicity of DarT*^Spu^* when expressed in *V. cholerae*, but Gp145^G62^ did not (Fig. 5C). These data are not consistent with the hypothesis that Gp145^G62^ functions as an antitoxin for another DarT homolog. However, the assays presented here were not meant to probe the full landscape of potential DarT-Gp145 interactions, and it remains possible that Gp145^G62^ could protect against DarT toxins not evaluated here.

## Discussion

To date, we have witnessed epidemic *V. cholerae* gain and lose genetic elements that confer defense against the prevalent vibriophage ICP1, and ICP1 gain and lose counter-defenses^15,20,25,29,50^. In this work, we characterize a new addition to this story—clinical *V. cholerae* isolates from 2008 to 2009 harbored a previously uncharacterized phage defense element that inhibits ICP1. We identify the root of anti-phage activity within the PDE as the toxin-antitoxin system DarTG. Before infection, DarT and DarG begin in a delicate balance that allows cells to grow without any defect. However, upon agitation by phage infection, the balance between the two is disrupted, and DarT, freed from DarG antagonism, halts phage genome replication (Fig. 2C), likely by adding ADP-ribose groups to ssDNA, which could sterically block phage DNA polymerase^31^. What precisely happens to the balance between DarT and DarG during phage infection has yet to be determined^31^. TA system activation following phage infection has been shown to occur in two ways: (1) phage-mediated host transcriptional and translational shutdown leads to degradation of the labile anitoxin^8^ or (2) a phage-encoded protein directly binds to a fused toxin-antitoxin, causing a conformational shift that renders the antitoxin inert^10^. Given these precedences, we hypothesize that activation of DarTG is dependent on DarG. If ICP1-mediated shutdown of host transcription and translation leads to degradation of DarG, DarG would no longer abrogate DarT’s toxicity both sterically and enzymatically. Alternatively, perhaps an ICP1-encoded protein interacts with DarG preventing its interaction with DarT and/or its catalytic activity. Previous work shows that ICP1 replication initiates within 8 minutes post-infection^22^, thus we expect DarTG activation to occur before 8 minutes post-infection. We further characterized the ICP1-DarTG conflict by isolating ICP1 mutants that escaped DarTG-mediated inhibition. This led to the identification of an ICP1-encoded protein (Gp145), which we propose to be called AdfB (for anti-DarT factor B). AdfB is expressed prior to 8 minutes post-infection and reaches peak expression by 8 minutes^51^. Thus, by the time DarT is active, AdfB functions as an antitoxin, abrogating DarT toxicity likely through direct interactions (Fig. 4), allowing ICP1 to produce progeny phage (Fig. 6).

**Figure 6:**
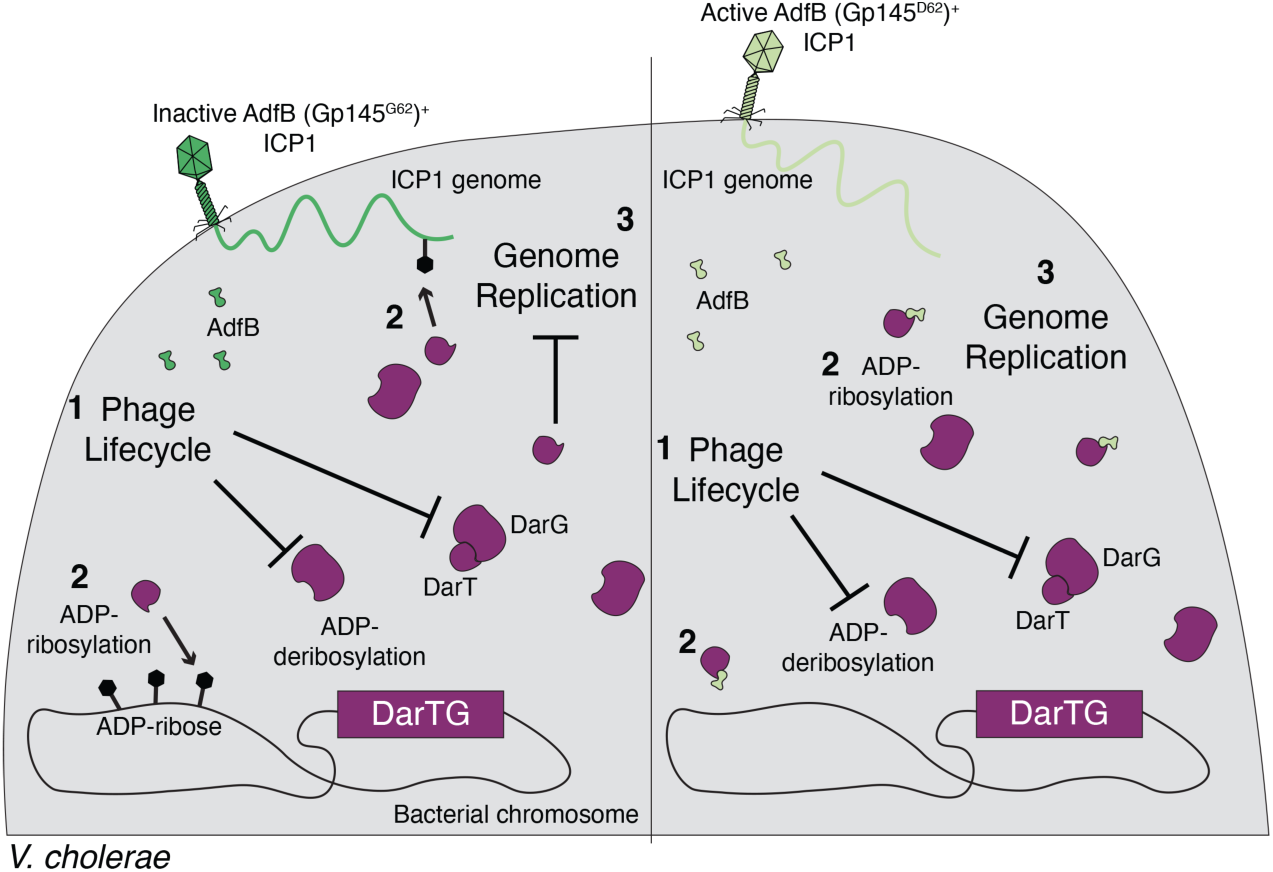
Proposed model of defense and counter-defense between DarTG^+^ *V. cholerae* and ICP1. (Left) When *V. cholerae* encodes DarTG, infection by ICP1 encoding the inactive allele of AdfB (Gp145^G62^) perturbs the balance between DarT and DarG by disrupting DarT-DarG binding and/or inhibiting DarG ADP-deribosylation (1). Now liberated from DarG, DarT ADP-ribosylates phage and bacterial genomic DNA (2), which halts ICP1 genome replication (3). (Right) When DarTG^+^ *V. cholerae* is infected by an ICP1 phage that encodes the active allele of AdfB (Gp145^D62^), the infection is still expected to perturb the balance between DarT and DarG (1). However, the now liberated DarT is antagonized by AdfB, inhibiting ADP-ribosylation (2), thus allowing ICP1 genome replication to continue (3).

There is an ongoing debate about the role of TA systems in phage defense. Our work further cements the DarTG system as a phage-defensive immune system. For the first time, we observe a DarTG system in clinical *V. cholerae* inhibiting the lytic phage, ICP1 (Fig. 2). Additionally, our work shows that DarTG inhibits phage from its native genomic locus. Furthermore, we provide evidence of DarT’s involvement in the stabilization of the putative MGE that carries it (Fig. S3). Previous studies of DarTG were unable to assess such activity given that they worked solely with plasmid-based expression systems of DarTG or entirely *in vitro*^30–32^. Combined, the features of DarTG providing phage defense and limiting MGE loss suggest that the system is multifunctional. Given that DarTG is associated with mobile genetic elements and other phage defense systems^30,31^, perhaps it serves a dual purpose in phage defense and protection of the defense element that encodes it. In this way, it could potentially protect other defense systems somewhat similar to how the TA system CreTA safeguards associated CRISPR-Cas immune systems^52^.

Multiple phages have been shown to encode their own antitoxins as a counter-defense against toxin-antitoxin systems, though how these antitoxin mimics arise is uncertain^8,31,39,41^. TA system inhibitors such as Dmd encoded by phage T4^39^ and pseudo-ToxI encoded by phage øTE^41^ abrogate their cognate toxin’s activity through direct interactions. Pseudo-ToxI could have arisen from an acquisition event where øTE infected ToxIN-encoding bacteria and gained the RNA antitoxin, or alternatively, the RNA could have evolved independently. The RnlA and LsoA inhibitor Dmd, however, has no apparent similarity to the antitoxins RnlB and LsoB^39^ and seems to require only one binding site to RnlA or LsoA compared to the two necessary for RnlB and LsoB^53,54^. These differences in mechanism indicate that Dmd likely evolved autonomously^8,54^. Concerning DarTG, phage RB69 is hypothesized to encode a direct DarT inhibitor, referred to as AdfA, which also shares low sequence similarity to DarG^31^. Given that the now characterized AdfB (Gp145) shares low sequence identity with the DarG (whose role it must fill), it suggests that it too arose independently. Other phages studied in *E. coli* can overcome DarTG via mutations in the phage-encoded DNA polymerase^31^. These mutations likely allow the DNA polymerase to bypass ADP-ribose groups to continue genome replication. Of our collection of vibriophages, only ICP1 was substantially inhibited by DarTG (Fig. S4). This indicates that podoviruses ICP2 and ICP3 might not activate the DarTG system, their polymerases can naturally bypass ADP-ribose groups, or they encode their own anti-DarT factor. Notably, none of the ICP1 escape mutants had DNA polymerase mutations (Supplemental Table 1) and attempts to evolve ICP1 isolates without AdfB on DarTG proved unsuccessful. Taken together with the fact that ICP1 encodes AdfB to overcome DarT, we hypothesize that ICP1’s DNA polymerase cannot tolerate mutations to allow bypassing of ADP-ribose groups. Additionally, ICP1 carrying a DarT antitoxin implies that it may not be able to evolve to avoid disrupting the balance between DarT and DarG during infection. If DarTG responds to cues integral to the phage lifecycle, such as host transcriptional shutdown that leads to rapid DarG turnover, then ICP1 would be unable to overcome DarTG unless it encoded something to fill the role of the now absent DarG, analogous to how phages overcome the depletion of antitoxins in both the ToxIN, RnlAB, and LsoAB systems^39,41^.

Prior to this study, ICP1’s evolution with clinical *V. cholerae* isolates was hypothesized to be driven by PLEs and SXT ICEs^20,25^. The discovery of the PDE characterized in this study, and an ICP1-encoded counter-defense in its core genome indicates that other defense elements not detected in our current methods of surveillance also drive ICP1 evolution. Interestingly, strains of ICP1 carry an active or inactive AdfB. Similarly, <RB69 and closely related phages T4 and T6 carry either an active or inactive allele of AdfA. A question yet to be answered is the purpose of these inactive alleles. For ICP1, we maintain the hypothesis that AdfB^G62^ could potentially be an antitoxin for diverse DarTG systems, given that our study was limited to testing one other DarTG system that inhibited ICP1 infection (Fig. 5). Additionally, these nonfunctional alleles of Adf could serve as indicators for potential co-circulating TA systems in the natural environment of each respective phage. Perhaps, <RB69 had not been faced with DarTG homologs like those tested in LeRoux *et al.* (2022)^31^, but phages T4 and T6 had. For <RB69, the inactive allele of AdfA was poised to mutate to ensure protection much like what we observed with AdfB in ICP1. Additionally, ICP1’s large genome size^15^ indicates that it has the space to accommodate nonessential genes, like the inactive allele of AdfB, whose deletion had no impact on ICP1 plaquing when infecting PDE^-^ *V. cholerae* in laboratory conditions (Fig. 3). Curiously, this does not seem to be the case in all ICP1’s counter-defenses. Against the PLE, ICP1 can encode either a CRISPR-Cas system or a nuclease (Odn) that occupy the same genomic locus in different strains of ICP1^28^. By being able to always accommodate AdfB but not CRISPR-Cas or Odn is suggestive that ICP1 can house smaller-sized counter-defenses in its core genome but something larger like ICP1’s CRISPR-Cas system (which is 10 times larger than Odn) is subject to more rapid turnover in the absence of selection pressure.

What drives the flux of MGEs in *V. cholerae* has yet to be understood. Looking at the prevalence of SXT ICE *Vch*Ind6 across time^20^, there does not seem to be massive shifts around the period when the PDE was circulating^20^. PLE3, in contrast, only seemed to be detected in clinical samples from 2008 – 2009 before being overtaken by PLE1^26^. Two prevailing hypotheses could explain PLE fluctuation. First, PLE evolution is driven by ICP1 equipped with appropriate counter-adaptations to the co-circulating PLE. Second, because PLEs require ICP1 late-expressed structural proteins^23^ and ICP1 genome replication machinery^24^ to horizontally transduce, a defense system that specifically inhibits ICP1 prior to PLE transmission could lead to PLE loss from the *Vibrio* population. Given that the PDE’s DarTG system inhibits ICP1 genome replication, we hypothesize that PLE3 could have been lost from the *V. cholerae* population because it was unable to transduce in the face of the co-circulating PDE. Here, our work highlights the power of using *V. cholerae* and co-isolated ICP1 to understand the evolutionary dynamics between bacteria and phages. The bacteria-phage co-evolutionary arms race has an ever-growing cast that is intertwined. The novel insights obtained from this work further implicate TA systems in phage-host interactions and highlight how non-SXT ICE and non-PLE defense elements shape both ICP1’s evolution but also MGE flux.

## Materials and Methods

### Bacterial and phage growth conditions

All *V. cholerae* strains used in this study, except for clinical strains, are derivatives of E7946 (listed in Supplemental Table 3). All bacterial strains were grown on LB agar from 20% glycerol stocks stored at -80°C overnight and then grown to stationary phase (OD_600_ greater than 1) in liquid LB broth with aeration. Strains were back-diluted from stationary phase into fresh LB broth to OD_600_=0.05 and grown to the specifications indicated for each assay below. When necessary, media was supplemented with antibiotics at the following concentrations: kanamycin (75 μg/ml), spectinomycin (100 μg/ ml), streptomycin (100 μg/ml), chloramphenicol (*V. cholerae* on plates) (2.5 μg/ml), chloramphenicol (*V. cholerae* in liquid) (1.25 μg/ml), chloramphenicol (*E. coli*) (25 μg/ml), and carbenicillin (50 μg/mL).

High titer phage stocks were generated as follows: Confluent lysis plates were made via the soft overlay method supplemented with appropriate antibiotics on their respective host *V. cholerae* strain (Supplemental Table 3). These plates were covered with STE Buffer (1M NaCl, 200mM Tris-HCl, 100mM EDTA) and rocked to harvest phages overnight at 4°C. The overlay was then collected into 50mL conical tubes and centrifuged at 4°C and 10,000 x g for 20 minutes. After transfer to a new tube, the supernatant containing phages was concentrated by centrifugation at 26,000 x g for 90 minutes. Phage pellets were nutated into fresh STE with agitation overnight at 4°C and then chloroform treated. Chloroform was cleared by centrifugation at 5,000 x g for 15 minutes.

### Strain construction and cloning

*V. cholerae* mutants, including the chromosomal expression constructs, were made by splice by overlap extension (SOE) PCR and introduced by natural transformation as previously described^55^. These PCR products contained either a spectinomycin resistance marker or a kanamycin resistance marker flanked by frt recombinase sites assembled to 1kB up and downstream regions of homology to the *V. cholerae* chromosome. PDE^+^ E7946 was engineered by repeated natural transformations to introduce parts the PDE with an antibiotic resistance gene into its natural integration site in E7946. Co-transformation with a SOE PCR product to remove the antibiotic marker in the PDE and a SOE PCR product to insert a spectinomycin resistance marker in the *lacZ* locus of *V. cholerae* was used to finalize the assembly of the PDE. The *lacZ* marker was removed via natural transformation of wild type *lacZ* and subsequent blue-white screening to identify strains with a repaired locus. After all this genetic manipulation, the strain was whole genome sequenced to verify the PDE and the rest of the genome were as expected. Chromosomal edits of DarT (DarT^E166A^) were made similarly: during natural transformation, both a SOE PCR product containing the mutated *darT* allele and another SOE PCR product with homology to the *lacZ* locus of *V. cholerae* containing a kanamycin resistance marker were added to *V. cholerae*. Natural transformants were selected on LB agar plates supplemented with the appropriate antibiotic and colony purified twice before stocking at -80°C in 20% glycerol. Plasmid constructs (Supplemental Table 4) were assembled via Gibson Assembly or Golden Gate Assembly and transformed into *E. coli* XL1-Blues. After sequence verification, plasmids were miniprepped and electroporated into *E. coli* S17s, which were then mated with *V. cholerae*. Plasmids for the inducible expression of genes of interest are derivatives of pMMB67EH with a theophylline-inducible riboswitch (riboswitch E) as described previously^56^. For the expression of diverse DarTG systems (Supplemental Table 2), genes were synthesized by GenScript and cloned into the pMMB67EH derivative.

Phage gene deletions and reconstitutions were constructed as previously described^57^. Here, an editing template with the desired deletion or point mutation was cloned into a plasmid and introduced to *V. cholerae*. This strain was infected with the ICP1 phage of interest. The multiple plaques that resulted from this plaque assay (described below) were picked into STE Buffer. ICP1 mutants were then used to infect *V. cholerae* with an induced Type 1-E CRISPR-Cas system containing a plasmid with a spacer targeting the phage gene of interest. Phages were purified three times on the targeting host before high titer stocks were made.

All deletions*, V. cholerae* mutants, engineered ICP1, and plasmid constructs were confirmed by Sanger sequencing.

### Phage plaquing assays

Spot plates and soft agar overlay were performed as previously described^56^. *V. cholerae* strains were grown to mid-log phase at 37°C with aeration. For spot plates, *V. cholerae* was added to 0.5% molten LB agar, vortexed, poured on a solid agar plate, and allowed to solidify. Once solidified, 3μL spots of 10-fold serial dilutions of phages were added and allowed to dry. These plates were incubated at 37°C overnight and imaged. For plaque assays to calculate the efficiency of plaquing (EOP), mid-log *V. cholerae* was mixed with pre-diluted phage samples. Phages were allowed to adsorb for 7 to 10 minutes before being added to 0.5% molten LB agar in individual petri dishes. These plates were incubated overnight at 37°C and plaques were counted. EOP was calculated by comparing the number of plaques formed on permissive *V. cholerae* relative to the number of plaques formed on a strain of interest.

If *V. cholerae* host strains contained plasmids, they were treated similarly as above, except antibiotics were supplemented as appropriate. For plasmids expressing DarTG*^Vch^*^A^and its homologs, strains were grown to OD_600_∼0.2 and induced with 1mM isopropyl β-D-1-thiogalactopyranoside (IPTG) and 1.5mM theophylline for 20 minutes. No induction was maintained in the petri dishes, but antibiotics were. For plasmids expressing spacers against genes of interest, strains were grown to OD_600_∼0.2 and induced with 1mM IPTG. Again, no inducer was maintained in the petri dishes, but appropriate antibiotics were.

### Identification of PDE-like elements

A BLASTn search was seeded with the nucleic acid sequence of the entire PDE. For initial insight into elements similar to the PDE, the BLASTn search was limited to *V. cholerae* strains in the NCBI database; however, only three non-redundant, annotated hits were selected from this search. To expand our search criteria, we then performed the search against the entire NCBI database. Here, the top 5 non-redundant, annotated hits were selected from NCBI, and associated nucleotide sequences were obtained. In both searches, any hit with a greater than 90% nucleotide identity and 75% query coverage were considered identical. Gene graphs of areas of homology between the PDE and other PDE-like elements and surrounding areas were generated using RStudio and the gggenes and ggplot packages. Genes predicted to be involved in phage defense were found by searching Defense Finder^4,58^ with the nucleic acid sequence of each predicted phage defense island.

### Transformation efficiency assay

Natural transformations to make deletions were conducted as described above. In addition to plating a serial dilution of competent *V. cholerae* exposed to SOE PCR product on LB agar plates containing the antibiotic for selection of the resistance marker, a serial dilution of competent bacteria was also plated on LB agar plates without antibiotic. Colony-forming units were enumerated on both sets of plates and the percentage of transformants were calculated by dividing the total number of transformants by the total number of *V. cholerae* colonies and multiplying by 100.

### Growth assays

*V. cholerae* strains expressing either DarT*^VchA^*and DarT*^VchA^*^E166A^ or DarT*^VchA/Spu^* and a vector expressing nothing, the DarT’s cognate DarG, Gp145^G62^, or Gp145^D62^ were back-diluted from stationary phase into fresh LB broth to OD_600_=0.05. Strains were grown to an OD_600_=0.3 at 37°C with aeration. A small aliquot was taken, serially diluted, and plated on LB agar plates. The remainder of the culture was induced with 0.01% arabinose and 1.5 mM theophylline and grown at 37°C with aeration for 3 hours. Strains with plasmids were also induced with 1mM IPTG. After growth, the cultures were serially diluted and plated on LB agar plates supplemented with antibiotics but not inducer. Plates were incubated at 37°C overnight, and colonies were counted the following day. Percent growth was calculated by dividing the number of colonies counted after three hours of growth by the number of colonies prior to induction. Antibiotics were included in the LB broth if strains contained plasmids.

### qPCR

Fold change in phage genome copy number was performed as previously described^24^. Briefly, *V. cholerae* strains were grown to an OD_600_=0.3 at 37°C with aeration and infected with ICP1 at a multiplicity of infection (MOI) of 0.1. A sample was taken immediately following the addition of phage to serve as the starting value and was boiled for 10 minutes. Infected cultures were incubated at 37°C with aeration for 20 minutes before the second sample was taken and boiled for 10 minutes. Boiled samples were diluted 1:50 to serve as the template for qPCR. All samples were run in biological quadruplet with technical replicates. Fold change was calculated as the amount of DNA in the sample at 20 minutes post-infection relative to the amount of DNA immediately following the addition of phage.

### ICP1 mutant selection and whole genome sequencing

PDE^+^ *V. cholerae* isolate 4 was used as a host for ICP1 in plaque assays as described above. Rare plaques were picked from biological replicates of this plaque assay and purified on PDE^+^ *V. cholerae* isolate 4 twice before a high-titer stock was generated. Phage stocks were treated with DNAase for 30 minutes at 37°C before heat inactivation at 65°C for 10 minutes. gDNA was collected using a Qiagen DNeasy blood and tissue DNA purification kit according to the manufacturer’s protocols. DNA libraries were prepared, and Illumina 150 x 150bp sequencing was performed by the Microbial Genome Sequencing Center (Pittsburgh, PA). The resulting genomes were assembled by SPAdes and analyzed by BreSeq (v0.33).

### Bacterial-two hybrid assays

Genes of interest were cloned into the multiple cloning sites of pUT18 and pKT25^49^ via Gibson Assembly. Specifically, we cloned a translational fusion between the C-terminus of the T25-CyaA catalytic domain and the catalytically inactive DarT^E166A^ (pT25C-*darT^E^*^166^*^A^*) and a translational fusion between the C-terminus of the T18-CyaA catalytic domain and both alleles of Gp145 (pT18C-*gp145^G^*^62^ and pT18C-*gp145^D^*^62^). Plasmids were electroporated into *E. coli* BTH101 in two biological replicates. We included the empty vector of plasmids pT18-CyaA and pT25-CyaA without inserts as negative controls. Three distinct colonies (two from the first electroporation and one from the second) were then picked into LB broth and grown at 37°C with aeration overnight. 10μL of this overnight culture was then spotted onto LB agar plates with kanamycin, carbenicillin, X-gal, and IPTG. Plates were incubated overnight at 30°C and then imaged.

### Phylogenetic analysis

All amino acid sequences used for the phylogenetic analysis are listed in Supplemental Data Sheet 2. The approximately-maximum likelihood phylogenetic trees comparing DarT alleles were conducted with FastTree (v2.1.10)^59^ using default settings. DarT homologs were gathered using two iterations of PSI-BLAST, where the input was the amino acid sequence of DarT*^Vch^*^A^. Sequences shorter than 100 and larger than 450 amino acids were omitted, and DarT sequences previously published^31^ were added if not present. 80% amino acid identity served as the cut-off for the clustering of branches. Additional visualization and annotation of the resulting files were performed with the iTOL (v6) web browser^60^.

## Supporting information

Supplementary Data Sheet 1

Supplementary Data Sheet 2

## Acknowledgments

The authors of this study would like to thank past and present members of the Seed Lab for fruitful discussion regarding this project. We would like to thank Reid T. Oshiro, Stephanie G. Hays, and Yamini Mathur for draft revisions and Drew Dunham for assistance with the bioinformatics. This project was supported by Grant number R01AI153303 to K.D.S. from the National Institute of Allergy and Infectious Diseases and its contents are solely the responsibility of the authors and do not necessarily represent the official views of the National Institute of Allergy and Infectious Diseases or NIH. K.D.S. holds an Investigators in the Pathogenesis of Infectious Disease Award from the Burroughs Wellcome Fund.

## Notes

### Competing Interest Statement

The authors have declared no competing interest.

