## Supplemental Figures 1-10 and Tables 1-4 for "Sporadic phage defense in epidemic *Vibrio cholerae* mediated by the toxin-antitoxin system DarTG is countered by a phage-encoded antitoxin mimic"

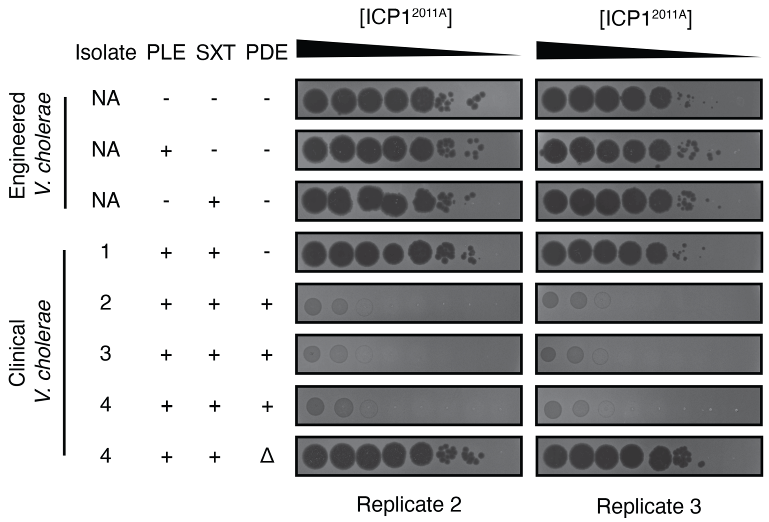


**Figure S1:** Biological replicates of ICP1$^{2011A}$ spotted on *V. cholerae* harboring different phage defenses.

Ten-fold serial dilutions of ICP1$^{2011A}$ were spotted on engineered or clinical *V. cholerae* strains containing PLE3, SXT *Vch*Ind6, or the PDE. Black zones of clearings are plaques. The opaque background is the *V. cholerae* lawn. Engineered strains are isogenic, while clinical isolates are not.

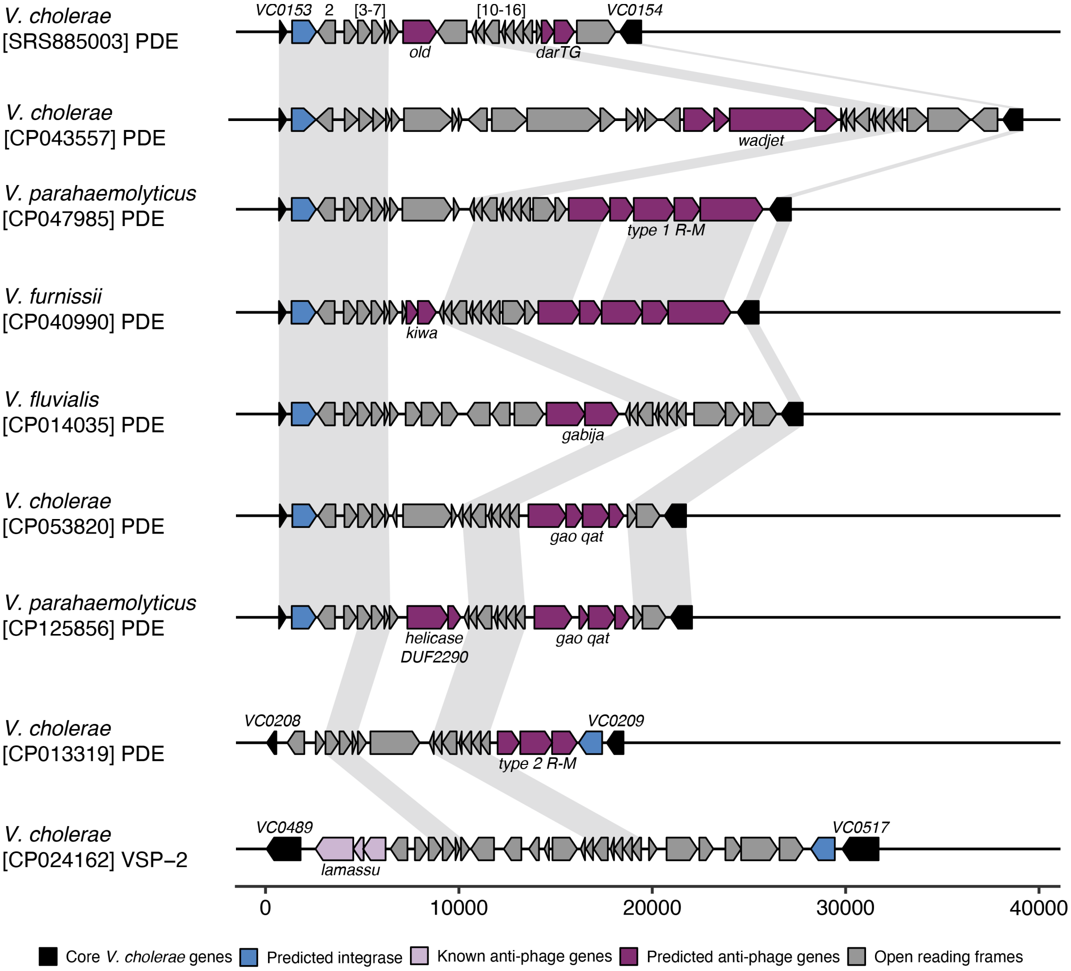


**Figure S2:** Elements related to the PDE also have putative anti-phage defense systems. Genetic maps (drawn to scale) of elements sharing nucleotide identify with the PDE characterized in this study (top). Defense Finder was used to predict if genes were involved in phage defense. Areas of >70% nucleotide identity are shaded in light grey. Accession numbers with the genomes containing these elements are written under the species name for each.

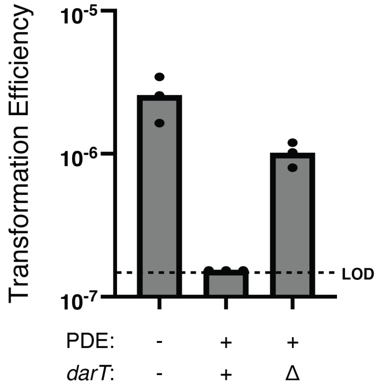


**Figure S3:** DarT limits natural transformation to engineer the deletion of the phage defense element (PDE) *in V. cholerae*.

Transformation efficiency (Transformants/total CFU) of isogenic PDE$^{-/+}$ *V. cholerae* and its derivatives with a PCR product with arms of homology to integrate a kanamycin resistance marker in place of the PDE (or in the genomic position of the PDE in PDE$^{-}$ *V. cholerae*). Each data point is representative of a single biological replicate, while the bar represents the mean of the replicates. LOD represents the limit of detection for the assay.

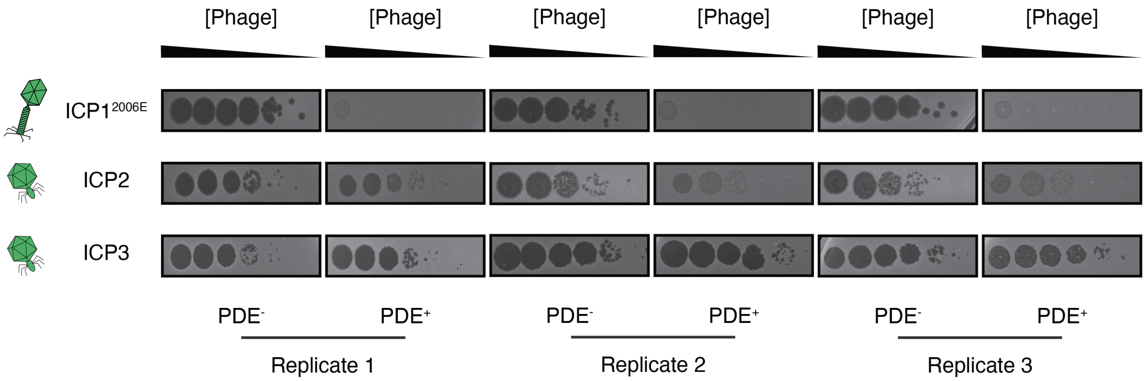


**Figure S4:** The PDE is sufficient to inhibit ICP1, but not all vibriophages. Ten-fold serial dilutions of ICP1$^{2006E}$, ICP2, and ICP3 were spotted on otherwise isogenic *V. cholerae* E7946 strains with and without the phage defense element (PDE). Black clearings are plaques where ICP1, ICP2, or ICP3 mounted a successful infection. The opaque background is the *V. cholerae* lawn.

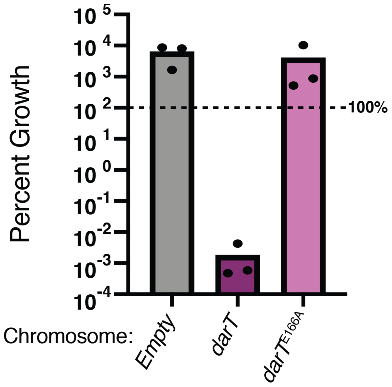


**Figure S5:** DarT$^{Vch\text{A E166A}}$ is not toxic to *V. cholerae.*
Percent growth as determined by colony-forming units of *V. cholerae* cells expressing the empty construct control, wild type DarT$^{Vch\text{A}}$, or DarT$^{Vch\text{A E166A}}$ via a chromosomal expression system for three hours. Each dot represents one biological replicate. The bar indicates the mean of all replicates. 100% cell survival is marked by the dotted line.

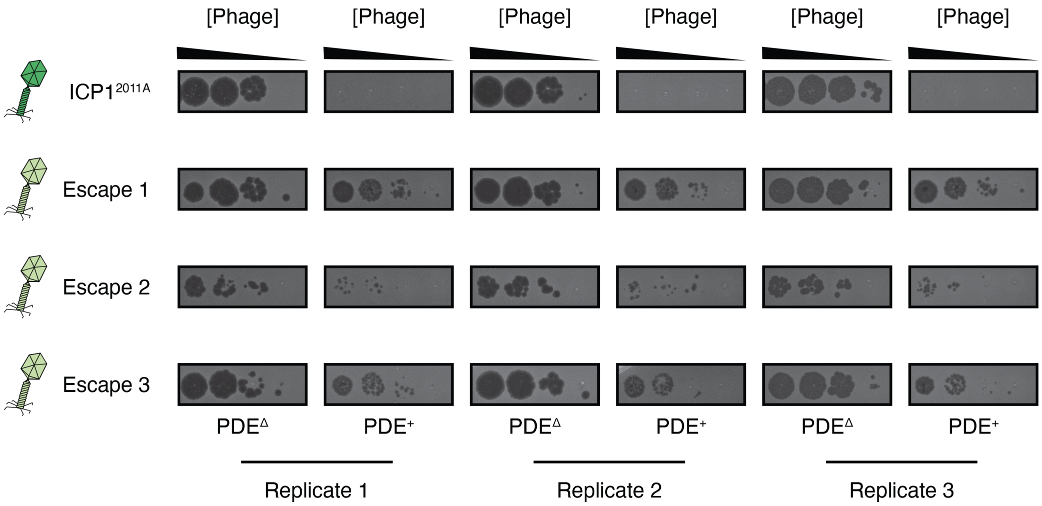


**Figure S6:** ICP1 escape mutants show improved EOP on PDE$^+$ clinical *V. cholerae.* Ten-fold serial dilutions of ICP1$^{2011A}$ and escape phage were spotted onto clinical *V. cholerae* strains encoding or lacking the phage defense element (PDE). Black clearings are where ICP1$^{2011A}$ was able to produce plaques. The opaque background is the *V. cholerae* lawn. Biological replicates are labeled.

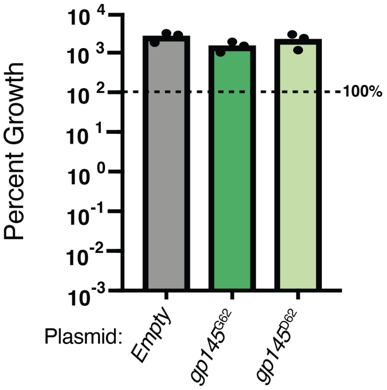


**Figure S7:** Neither allele of Gp145 induces DarTG-mediated cell death.
Percent growth as determined by colony-forming units of PDE+ *V. cholerae* cells expressing an empty vector, Gp145$^{G62}$, or the evolved Gp145$^{D62}$ variant for three hours. Each dot represents one biological replicate. The bar indicates the mean of all replicates. 100% cell survival is marked by the dotted line.

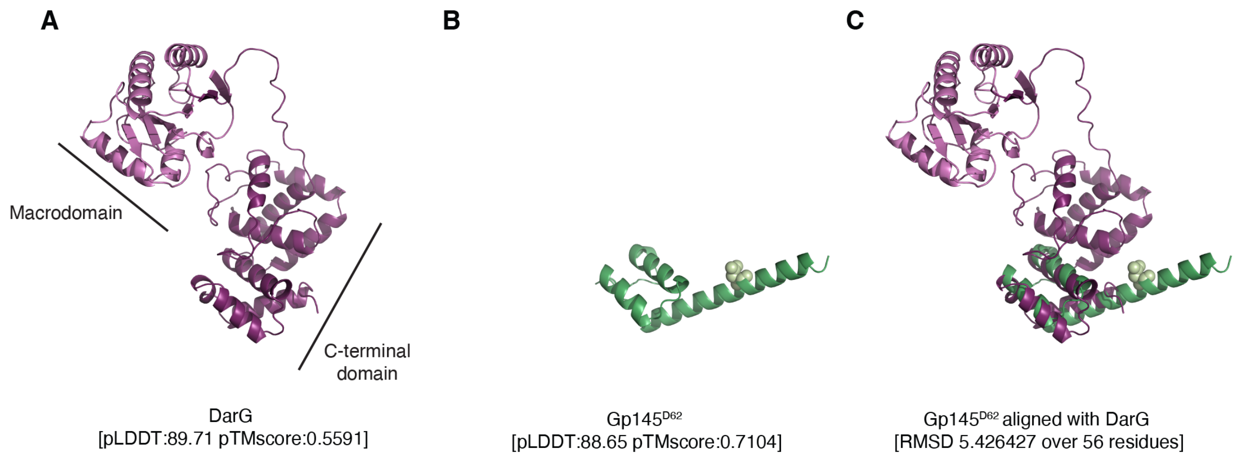


**Figure S8:** Gp145$^{D62}$ and the C-terminus of DarG are predicted to be structurally similar.

A) The AlphaFold2[48] predicted structure of DarG reveals two major domains. The light purple domain (N-terminus domain) is the macrodomain, known for ADP-deriboyslation. The dark purple domain is the C-terminus, which is known to bind to DarT in other systems[38]. pLDDT and pTM scores are written below the structure.

B) AlphaFold2 predicted structure of Gp145$^{D62}$ is shown in green. The D62 residue is shown in spheres and a lighter green color. pLDDT and pTM scores are written below the structure.

C) The superimposition of predicted structures of Gp145$^{D62}$ and DarG reveals that the C-terminus of DarG and Gp145$^{D62}$ show a striking structural similarity. The RMSD value is shown below the structure.

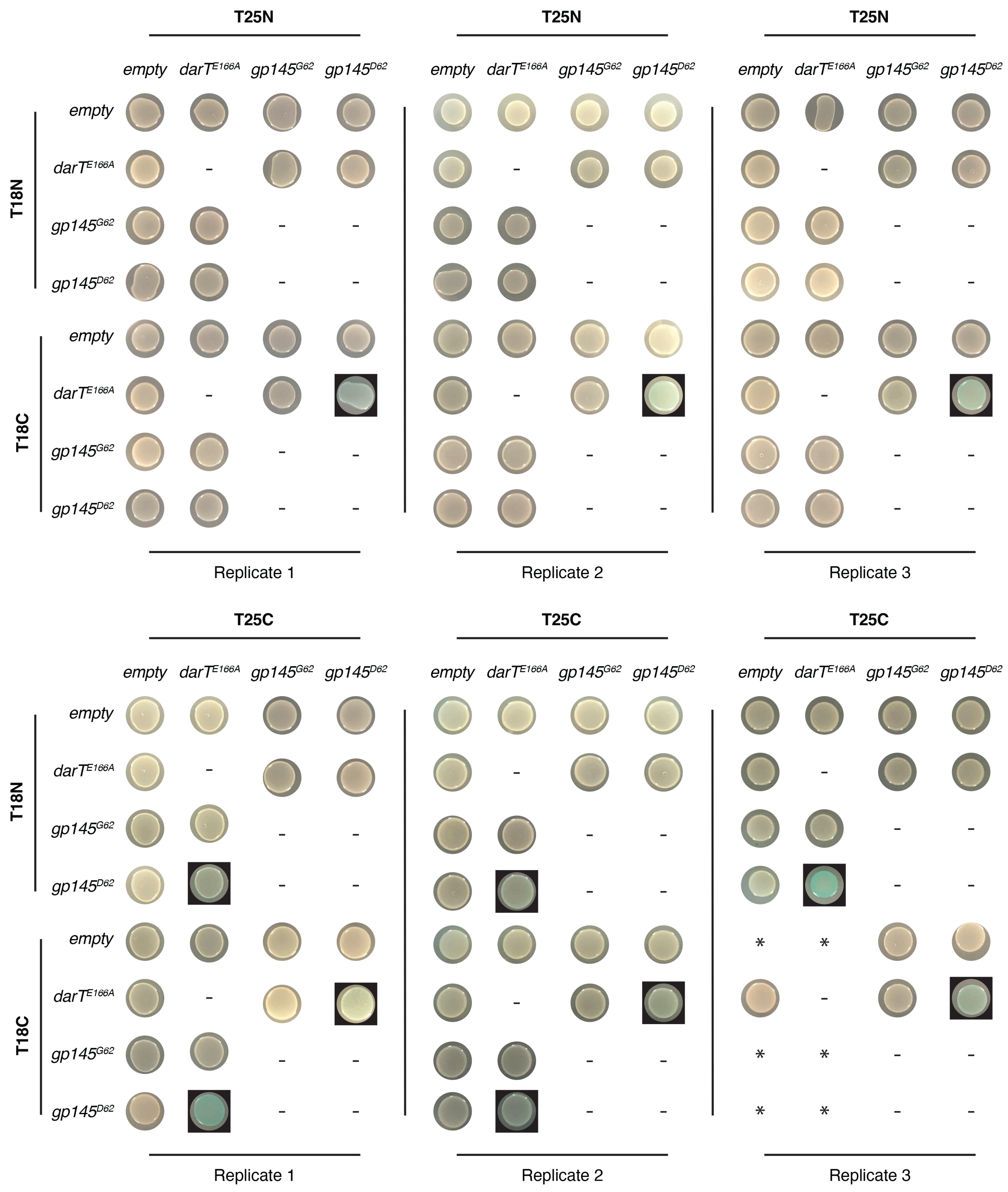
T25N
T25N
T25N
empty
darT^E166A
gp145^G62
gp145^D62
T18N
T18C
Replicate 1
Replicate 2
Replicate 3
T25C
T25C
T25C
*

**Figure S9:** Replicates of bacterial two-hybrid data to detect protein-protein interactions between both alleles of Gp145 and DarT$^{E166A}$.

Cells containing plasmids indicated on the top and left were spotted on agar plates containing X-gal and inducer. Blue spots indicate a physical interaction between the proteins fused to the CyaA subunits. Any blue colony is shown highlighted with a black box. Biological replicates are labeled. Any combination with a “-” were not tested, while the combinations with an “*” are shown in the main text.

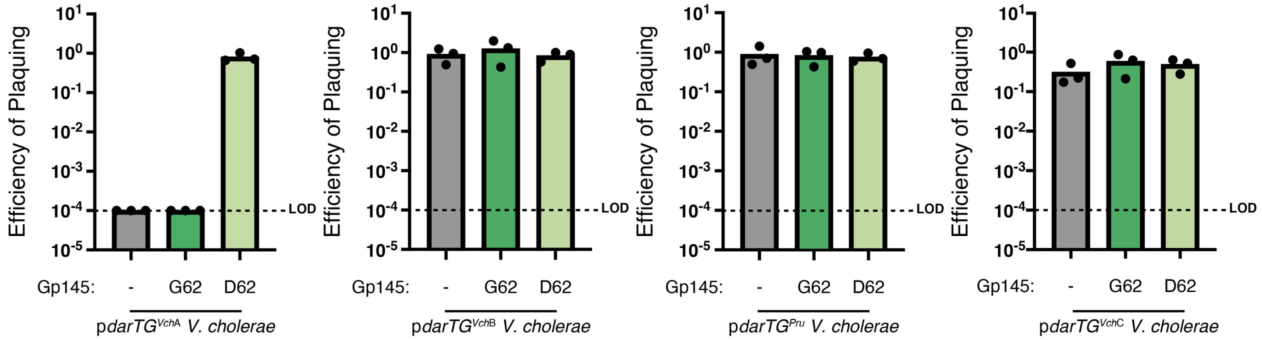


**Figure S10:** Diverse DarTG systems do not inhibit ICP1 lacking Gp145 or expressing either Gp145$^{G62}$ or Gp145$^{D62}$.

The efficiency of plaquing (EOP) of engineered ICP1$^{2006E}$ with the allele of *gp145* indicated on *V. cholerae* expressing different DarTG systems from a plasmid (additional information for each DarTG system is in Supplemental Table 2). Each data point is representative of a single biological replicate, while the bar represents the mean of these replicates. LOD is the limit of detection.

Supplemental Table 1: BreSeq analysis of ICP1$^{2011A}$ escape mutants selected on PDE$^{+}$ clinical *V. cholerae*.

| Position | Escape 1[1] | Escape 2 | Escape 3 | Annotation | Gene[2] |
|---|---|---|---|---|---|
| 14,612 | | | 6.00% | F72S (TTT→TCT) | Gp37 ← |
| 17,642 | 100% | | | intergenic (-202/-36) | Gp46 ← / → Gp47 |
| 73,244 | 100% | | | E142G (GAG→GGG) | Gp124 ← |
| 76,467 | | | 7.30% | F82Y (TTT→TAT) | Gp127 ← |
| 76,470 | | | 7.10% | K81T (AAG→ACG) | Gp127 ← |
| 76,473 | | | 7.00% | R80L (CGT→CTT) | Gp127 ← |
| 76,476 | | | 7.40% | I79K (ATA→AAA) | Gp127 ← |
| 80,258 | | | 5.60% | K56E (AAG→GAG) | Gp131 ← |
| **86,240** | **100%** | **100%** | **100%** | **G62D (GGT→GAT)** | **Gp145 ←** |
| 112,450 | 100% | | | coding (1807-1851/1917 nt) | Gp203 → |
| [1]Frequency scores are the percentage of reads with the mutation compared to the total reads. | | | | | |
| [2]Arrows represent the direction of the strand. Arrows to the left indicate 3' to 5'. Arrows to the right indicate 5' to 3'. | | | | | |

Supplemental Table 2: Summary of DarTG systems used in this study.

| Name | Species | DarT NCBI Reference Sequence | DarT Percent Identity to DarT$^{VchA}$ | DarG NCBI Reference Sequence | DarG Percent Identity to DarG$^{VchA}$ |
|---|---|---|---|---|---|
| DarTG$^{VchA}$ | *Vibrio cholerae* | See data sheet 1 for Amino Acid Sequence | 100 | See data sheet 1 for Amino Acid Sequence | 100 |
| DarTG$^{Spu}$ | *Shewanella putrefaciens* | WP_011920083.1 | 80.84 | WP_011920084.1 | 86.71 |
| DarTG$^{VchB}$ | *Vibrio cholerae* | WP_198302751.1 | 62.62 | WP_096070761.1 | 79.83 |
| DarTG$^{PruA}$ | *Providencia rustigianni* | WP_006813742.1 | 65.89 | WP_039854900.1 | 77.52 |
| DarTG$^{VchC}$ | *Vibrio cholerae* | WP_025795243.1 | 13.42 | WP_057552482.1 | 24.31 |

Supplemental Table 3: Strains used in this study.
*Numbers refer to the reference listed in the main text. References with roman numerals in parenthesis are additional references listed below the tables.

| Name in text | Strain | Description | Source* |
|---|---|---|---|
| Permissive PDE$^{-}$ *V. cholerae* | KDS6 | *Vibrio cholerae* E7946, El Tor Ogawa, O1, streptomycin resistant | Levine et al., 1982[(I)] |
| PLE3$^{+}$ engineered *V. cholerae* | KDS38 | *Vibrio cholerae* E7946 containing PLE3 integrated between VCA0415 and VCA0416. | O'Hara et al., 2017[25] |
| SXT ICE$^{+}$ engineered *V. cholerae* | KL550 | *Vibrio cholerae* E7946 containing SXT ICE *Vch*Ind6 | LeGault et al., 2022[20] |
| Clinical *V. cholerae* isolate 1 | KS511 | *Vibrio cholerae* O1 clinical isolate from 2008 [SRR1944526] | Dalia et al., 2014[55] |
| Clinical *V. cholerae* isolate 2 | KS515 | *Vibrio cholerae* O1 clinical isolate from 2009 [SRR1944520] | Dalia et al., 2014[55] |
| Clinical *V. cholerae* isolate 3 | KS516 | *Vibrio cholerae* O1 clinical isolate from 2009 [SRR1944521] | Dalia et al., 2014[55] |
| Clinical *V. cholerae* isolate 4 | KS39 | *Vibrio cholerae* O1 clinical isolate from 2009 [SRR1944525] | Dalia et al., 2014[55] |
| Clinical *V. cholerae* isolate 4 ΔPDE | KMP1049 | KS39 with an in-frame frt cassette replacing *orfs1-20* of the PDE | This study |
| Clinical *V. cholerae* isolate 4 Δ*darT* | KMP994 | KS39 with an in-frame frt cassette replacing *darT* of the PDE | This study |
| PDE$^{+}$ *V. cholerae* | KMP268 | KDS6 containing the PDE integrated between VC0153 and VC0154 | This study |
| PDE$^{+}$ *V. cholerae* Δ*old* | KMP298 | KMP268 with an in-frame frt cassette replacing *old* | This study |
| PDE$^{+}$ *V. cholerae* Δ*darT* | KMP375 | KMP268 with an in-frame frt cassette replacing *darT* | This study |

| | | | |
|---|---|---|---|
| PDE+ *V. cholerae* *darT*$^{E166A}$ | KMP770 | KMP268 (with an in-frame codon replacement (GAG->GCG) to have *DarT*$^{E166A}$ expressed, *lacZ* replaced with Kanamycin resistance cassette integrated into *V. cholerae lacZ* locus | This study |
| PDE- *V. cholerae* chromosomal expression empty | KMP594 | KDS6 with expression cassette ($P_{BAD}$-riboswitchE) from Dalia *et al*., 2020[II] integrated into *V. cholerae* lacZ locus with Kanamycin resistance cassette, expresses nothing (empty) | This study |
| PDE- *V. cholerae* chromosomal expression *darT*$^{VchA+}$ | KMP596 | KMP594 with DarT$^{VchA}$ expressed from the expression cassette | This study |
| PDE- *V. cholerae* chromosomal expression *darT*$^{VchA\ E166A+}$ | KMP592 | KMP594 with DarT$^{VchA\ E166A}$ expressed from the expression cassette | This study |
| Empty Vector+ PDE- *V. cholerae* chromosomal expression *darT*$^{VchA+}$ | KMP627 | KMP596 containing an empty vector | This study |
| p*darG*$^{VchA+}$ PDE- *V. cholerae* chromosomal expression *darT*$^{VchA+}$ | KMP630 | KMP596 containing a vector that can express DarG$^{VchA}$ | This study |
| p*gp145*$^{G62+}$ PDE- *V. cholerae* chromosomal expression *darT*$^{VchA+}$ | KMP643 | KMP596 containing a vector that can express Gp145$^{G62}$ | This study |
| p*gp145*$^{D62+}$ PDE- *V. cholerae* chromosomal expression *darT*$^{VchA+}$ | KMP631 | KMP596 containing a vector that can express Gp145$^{D62}$ | This study |

| | | | |
|---|---|---|---|
| PDE⁻ *V. cholerae* chromosomal expression *darT*$^{Spu+}$ | KMP1193 | KMP594 with DarT$^{Spu}$ expressed from the expression cassette | This study |
| Empty Vector⁺ PDE⁻ *V. cholerae* chromosomal expression *darT*$^{Spu+}$ | KMP1195 | KMP1193 containing an empty vector | This study |
| p*darG*$^{Spu+}$ PDE⁻ *V. cholerae* chromosomal expression *darT*$^{Spu+}$ | KMP1201 | KMP1193 containing a vector that can express DarG$^{SpuA}$ | This study |
| p*gp145*$^{G62+}$ PDE⁻ *V. cholerae* chromosomal expression *darT*$^{Spu+}$ | KMP1197 | KMP1193 containing a vector that can express Gp145$^{G62}$ | This study |
| p*gp145*$^{D62+}$ PDE⁻ *V. cholerae* chromosomal expression *darT*$^{Spu+}$ | KMP1199 | KMP1195 containing a vector that can express Gp145$^{D62}$ | This study |
| BACTH host | KDS179 | *E. coli* without adenylate cyclase gene used for BACTH | McKitterick and Seed, 2018[56] |
| ICP1$^{2011A}$ | KSϕ40 | ICP1_2011_Dha_A WT Accession Number: MH310933.1 | Seed et al., 2013[29] |
| ICP1$^{2006E}$ | KSϕ36 | ICP1_2006_Dha_E (Accession Number: MH310934.1) Δ*CRISPR* Δ*cas2-3* | McKitterick and Seed, 2018[56] |
| ICP2 | ICP2 | Accession Number: HQ641345 | Seed et al., 2011[15] |
| ICP3 | ICP3 | Accession Number: HQ641340 | Seed et al., 2011[15] |
| ICP1$^{2006E}$ Δ*gp145* | KMPϕ34 | KSΦ36 containing an in-frame deletion of *gp145* | This study |
| ICP1$^{2006E}$ *gp145*$^{D62}$ | KMPϕ38 | KMPΦ34 where *gp145* locus has been repaired with *gp145*$^{D62}$ and a 3X FLAG tag on the C-terminus | This study |

| | | | |
|---|---|---|---|
| ICP1$^{2006E}$ *gp145*$^{G62}$ | KMPϕ40 | KMPΦ34 where *gp145* locus has been repaired g*p145*$^{G62}$ a 3X FLAG tag on the C-terminus | This study |
| ICP1$^{2011A}$ Escape 1 | KMPϕ6 | KSΦ36 that now possesses a mutation in Gp145, now codes for Gp145$^{D62}$, and possesses intergenic mutations and Gp124 mutation (described in Supplemental Table 1) | This study |
| ICP1$^{2011A}$ Escape 2 | KMPϕ9 | KSΦ36 that now possesses a mutation in Gp145, now codes for Gp145$^{D62}$ | This study |
| ICP1$^{2011A}$ Escape 3 | KMPϕ12 | KSΦ36 that now possesses a mutation in Gp145, now codes for Gp145$^{D62}$ | This study |

86

Supplemental Table 4: Plasmids used in this study.

| **Plasmids** | **Description and Identifier** | **Source** |
|---|---|---|
| Empty Vector | pMMB67EH engineered to contain an inducible riboswitch (E- induced by theophylline) downstream of a $P_{tac}$ promoter, KS1864 | Lab collection |
| p*darTG*$^{VchA}$ | Empty vector engineered for the inducible expression of DarTG$^{VchA}$, KMP416 | This study |
| p*darT*$^{E166A}$*G*$^{VchA}$ | Empty vector engineered for the inducible expression of DarT$^{E166A}$G$^{VchA}$, KMP780 | This study |
| p*gp145*$^{G62}$ | Empty vector engineered for the inducible expression of Gp145$^{G62}$, KMP418 | This study |
| p*gp145*$^{D62}$ | Empty vector engineered for the inducible expression of Gp145$^{D62}$, KMP420 | This study |
| p*darG*$^{VchA}$ | Empty vector engineered for the inducible expression of DarG$^{VchA}$, KMP518 | This study |
| *empty* pT18 | pUT18, T18 subunit of *cya* | Lab collection |
| *empty* pT25 | pKNT25, T25 subunit of *cya* | Lab collection |
| p*darT*$^{E166A}$ - T25N | pKNT25, T25 subunit of *cya* fused to N-terminus of *darT*$^{E166A}$, KMP701 | This study |
| p*gp145*$^{G62}$- T25N | pKNT25, T25 subunit of *cya* fused to N-terminus of *gp145*$^{G62}$, KMP705 | This study |
| p*gp145*$^{D62}$- T25N | pKNT25, T25 subunit of *cya* fused to N-terminus of *gp145*$^{D62}$, KMP707 | This study |
| p*darT*$^{E166A}$ - T18N | pUT18, T18 subunit of *cya* fused to N-terminus of *darT*$^{E166A}$, KMP717 | This study |
| p*gp145*$^{G62}$- T18N | pUT18, T18 subunit of *cya* fused to N-terminus of *gp145*$^{G62}$, KMP721 | This study |
| p*gp145*$^{D62}$- T18N | pUT18, T18 subunit of *cya* fused to N-terminus of *gp145*$^{D62}$, KMP723 | This study |
| p*darT*$^{E166A}$ - T25C | pKNT25, T25 subunit of *cya* fused to C-terminus of *darT*$^{E166A}$, KMP709 | This study |
| p*gp145*$^{G62}$- T25C | pKNT25, T25 subunit of *cya* fused to C-terminus of *gp145*$^{G62}$, KMP713 | This study |
| p*gp145*$^{D62}$- T25C | pKNT25, T25 subunit of *cya* fused to C-terminus of *gp145*$^{D62}$, KMP715 | This study |
| p*darT*$^{E166A}$ - T18C | pUT18, T18 subunit of *cya* fused to C-terminus of *darT*$^{E166A}$, KMP725 | This study |

| | | |
|---|---|---|
| p*gp145*$^{\mathrm{G62}}$-T18C | pUT18, T18 subunit of *cya* fused to C-terminus of *gp145*$^{\mathrm{G62}}$, KMP729 | This study |
| p*gp145*$^{\mathrm{D62}}$-T18C | pUT18, T18 subunit of *cya* fused to C-terminus of *gp145*$^{\mathrm{D62}}$, KMP731 | This study |
| p*darTG*$^{Spu}$ | Empty vector now engineered for the inducible expression of DarTG$^{Spu}$ downstream of promoter and riboswitch, KMP699 | This study |
| p*darG*$^{Spu}$ | Empty vector now engineered for the inducible expression of DarG$^{Spu}$ downstream of promoter and riboswitch, KMP749 | This study |
| p*darTG*$^{Vch\mathrm{B}}$ | Empty vector now engineered for the inducible expression of DarTG$^{Vch\mathrm{B}}$ downstream of promoter and riboswitch, KMP867 | This study |
| p*darTG*$^{Pru}$ | Empty vector now engineered for the inducible expression of DarTG$^{Pru}$ downstream of promoter and riboswitch, KMP697 | This study |
| p*darTG*$^{Vch\mathrm{C}}$ | Empty vector now engineered for the inducible expression of DarTG$^{Vch\mathrm{A}}$ downstream of promoter and riboswitch, KMP855 | This study |
